# Urbanization effects are trait-specific and city-dependent across a widespread spider’s global range

**DOI:** 10.64898/2026.06.19.733406

**Authors:** Bram Vanthournout, Maxime Dahirel, Angela Chuang, Katrien De Wolf, Nel D’haenekint, Isabel S. Abihssira-Garcia, Angela M Alicea-Serrano, Markus Andersen, Hannah Anderson, Leticia Aviles, James B Barnett, Elif Başarır, Thomas R Beatman, Jesper Bechsgaard, Trine Bilde, Camillo Elia Biundo, Jessica C. Boles, Erin E Brandt, Snata Chakraborty, Alix Coonfield, Solène Croci, Jordan P. Cuff, Mario Driesen, Sebastian Echeverri, Kirsten R Engeseth, Ignacio Escalante, Lauren A Esposito, Andreas Fischer, Saoirse Foley, Miriam Julia Frutiger, Reese Gartly, Laura Garzoli, Jacob Gorneau, Leanne Grieves, Jennifer Guevara, Andrea Haberkern, Camela Haddad, Fredrik Ø. Hanslin, Thomas Hesselberg, Chiara Hirschkorn, Emmanuel Hung, Marco Isaia, Morgan Jackson, Elizabeth Jakob, Warbota Khum, Jihoo Kim, Gitte Kragh, Simona Kralj-Fišer, Amalia Aikaterini Mailli, Stefano Mammola, Marika Maranzano, Nicolas Maughan, Sean McCann, Radek Michalko, Deepti Manjari Patel, Richard Pearce, Julien Pétillon, Elena Piano, Joost A. M. Raeymaekers, Daniela Christina Rößler, Indra Saenen, Orlando Schwery, Catherine Scott, Laini Taylor, Zoe Umbach, Martijn L. Vandegehuchte, Nathan Viel, Trinity Walls, Alyssa Wilkinson, Alex M Winsor, Shiyang Wu, Lin Yan, Eric Yip, Liliana D’Alba, Matthew Shawkey, Dries Bonte

## Abstract

Urban environments impose strong selective pressures through biotic and abiotic factors, driving changes in behavior, physiology, and morphology. Yet, responses vary across taxa and cities, and it remains unclear which traits respond consistently and what factors moderate this variation. We addressed these questions using the widespread European garden spider (*Araneus diadematus*) as a model, measuring size, color, and web-building traits along urban–rural transects in 22 cities across its distribution range. Using a meta-analytic framework, we assessed how city-specific characteristics influenced trait variation. Urbanization consistently reduced relative abdomen surface area, a proxy for body condition. Exploratory meta-regressions suggest that web-building response was predicted by temperature: compared to their non-urban surroundings, urban webs are larger in colder regions and smaller in warmer regions. In contrast, body size and abdomen brightness varied among cities without clear environmental predictors. These findings show that urbanization effects are trait- and context-dependent, likely influenced by local factors such as heat island intensity, microclimate, or prey availability. Linking within- and between-city variation will improve understanding of species phenotypic responses to urban environments.

## Introduction

Urbanization is a major driver of human-induced rapid environmental change. Over half of the global human population now lives in cities, and urban areas are projected to grow by 2.5 billion people by 2050, alongside substantial expansion of urban land (United Nations, 2019; Chen et al., 2020). These land-use changes create novel ecosystems with distinct selection pressures compared to semi-natural environments (Johnson & Munshi-South, 2017; Parris, 2016). Abiotic pressures include artificial light at night, noise pollution, chemical contaminants, impervious surfaces, elevated temperatures from urban heat islands, and altered hydrology (Alberti, 2015). The expansion of urban areas also results in extensive habitat loss and fragmentation, which disrupts dispersal patterns and influences gene flow and genetic drift (Alberti, 2015). At the community level, urban environments filter taxonomic and functional diversity, reducing species richness, especially in terrestrial arthropods (Fenoglio, Rossetti, & Videla, 2020) and homogenizing species assemblages (Aronson et al., 2016; Merckx & Van Dyck, 2019; Piano, Souffreau, et al., 2020; Piquet, Piano, Tolve, & Isaia, 2024). These ecological shifts alter species interactions (Miles, Breitbart, Wagner, & Johnson, 2019a; Murray et al., 2019; Theodorou, 2022), for example, by changing prey availability and quality (El-Sabaawi, 2018).

Species respond to urban environments through diverse morphological, physiological, and behavioral changes (Alberti et al., 2017), driven by genetic adaptations or phenotypic plasticity (Diamond & Martin, 2021; Lambert, Brans, Des Roches, Donihue, & Diamond, 2021; Thompson, Capilla-Lasheras, Dominoni, Réale, & Charmantier, 2022). Importantly, these responses are often taxon- and species-specific (Naidoo, Chamberlain, & Reynolds, 2024), with contrasting patterns, such as an increase, decrease or no response in body size in different arthropod groups (Kralj-Fišer et al., 2017, Merckx, Souffreau, et al., 2018, Van den Bossche et al., 2025). Moreover, intraspecific responses to urbanization are not always uniform and can vary substantially between cities (Diamond, Chick, Perez, Strickler, & Martin, 2018; Salmón et al., 2023; Santangelo et al., 2022).

An outstanding question in urban ecology is whether similar urban pressures drive parallel evolutionary or plastic responses (Verrelli et al., 2022). Syntheses highlight significant variation in responses, often due to studies focusing on single cities (Szulkin, Munshi-South, & Charmantier, 2020). However, while urban environments share broad similarities, cities differ markedly in characteristics that shape the direction and strength of selection. For example, city size influences ecologically relevant factors (Uchida et al., 2021) such as the intensity of the urban heat island effect, with more populous cities being warmer (Manoli et al., 2019). Additionally, city size might impact dispersal rates and, hence, alter gene flow with urban landscape features functioning as barriers or conduits (Richardson, Urban, Bolnick, & Skelly, 2014; Miles et al., 2019b). Contrast with the surrounding environment of a city and the presence of green spaces within a city can further determine the strength of urbanization effects. Moreover, the latitude and associated regional climate of a city can interact with these local factors, leading to variable responses (Diamond, Dunn, Frank, Haddad, & Martin, 2015; Lövei & Magura, 2022; Youngsteadt, Ernst, Dunn, & Frank, 2017).

Few studies have investigated how urban-rural clines of phenotypic traits are modulated by climate-dependent environmental factors across a sufficiently large number of cities and range of latitudes. Existing work suggests that temperature (Cosentino & Gibbs, 2022) and potential evapotranspiration coupled with vegetation cover (Santangelo et al., 2022) can moderate responses to urbanization, indicating that predictable, parallel evolutionary changes occur, though local conditions ultimately determine their magnitude and direction. To better understand these environmental, macro-ecological drivers and the extent of potential parallel responses in urban environments, more comprehensive studies are needed. Such investigations should ideally involve sampling across many cities with diverse characteristics and span a broad range of climatic conditions (Johnson & Munshi-South, 2017). A critical prerequisite for this approach is the selection of a study species that occurs in both urban and rural habitats and possesses a sufficiently wide geographic distribution.

Spiders are dominant arthropod predators in many terrestrial ecosystems and occupy a wide range of habitats (Foelix, 2010). As ectotherms, spider juvenile development (Li & Jackson, 1996; Thorbek, Sunderland, & Topping, 2003) and adult activity (Hesselberg & Vollrath, 2006) are also strongly influenced by environmental temperature. For web-building species, the structure of capture webs (capture area, mesh size) is a highly plastic trait, directly linked to individual foraging decisions influenced by environmental factors such as temperature and prey size (Vollrath, Downes, & Krackow, 1997). This physiological sensitivity and reliance on webs for prey capture makes web-building spiders well-suited for investigating the impacts of various urban stressors, such as altered prey availability and quality, as well as increased temperatures.

Spiders are increasingly used in urban ecology research (reviewed by Willmot et al. 2025), revealing responses to urbanization such as a reduced species abundance and diversity (Piano, Souffreau, et al., 2020), community homogenization (Piano, Giuliano, & Isaia, 2020), and responses to urban stressors including heat island effects (Clark & Johnson, 2024), artificial light at night, and altered prey dynamics. For example, artificial light at night can accelerate juvenile development and reduce body size (Willmott et al., 2018) while increasing prey capture (Gomes, 2020). Other studies report reduced light avoidance (Czaczkes et al., 2018), shifts in nutrient composition (Trubl & Johnson, 2019), and scale-dependent dispersal effects (Bonte et al., 2023). Morphological and behavioral responses vary widely, with body size and web dimensions showing inconsistent patterns across spider species and cities (Cabon et al., 2024; Dahirel et al., 2019; Lowe et al., 2016; Ripp et al., 2018). Some species even act as urban exploiters (Clark & Johnson, 2024; Lowe et al., 2016). Beyond size and web architecture, body coloration may also respond to urbanization. Potential drivers include altered camouflage needs, pollution-induced melanism (Goiran et al., 2017), resource limitations affecting pigmentation (Leveau, 2021; Salmón et al., 2023), and thermal melanism, where paler individuals absorb less heat in warmer urban environments (Rao & Mendoza-Cuenca, 2016; Stuart-Fox et al., 2017; Trullas et al., 2007).

This study investigates how urbanization affects ecologically consequential trait responses, and how they are moderated by city characteristics and environmental factors. We focus on the European garden spider (*Araneus diadematus* Clerck, 1757), a widely distributed orb-weaver (Araneidae) occurring across natural and anthropogenic habitats throughout the Holarctic. Native to the Palearctic and introduced to North America in the late 19th century (Dondale & Redner, 2003), *A. diadematus* constructs wheel-shaped orb webs that respond plastically to prey size with larger meshes and capture areas for larger prey (Schneider & Vollrath, 1998), and to environmental conditions, with smaller, rounder webs in windy sites and smaller meshes under lower temperatures in laboratory settings (Vollrath et al., 1997). Webs are typically rebuilt daily (Breed et al., 1964), providing a dynamic record of foraging investment. As a generalist predator, the species captures a wide range of arthropod prey (van Schrojenstein Lantman et al., 2021) and exhibits striking variation in body coloration, from light to dark hues of brown, orange, red, yellow, gray, and black (Messas et al., 2025).

*A. diadematus* exhibits phenotypic changes in response to both local and landscape-level urbanization. In Belgium, Dahirel et al. (2019) reported smaller urban spiders, consistent with predictions based on reduced prey biomass in cities (De Wolf et al., 2025) and increased metabolic rates due to higher urban heat island temperatures. Urban spiders also build smaller webs with smaller mesh sizes at a local scale of urbanization, but larger webs at landscape scale, likely reflecting a behavioural response to reduced prey sizes in urban areas. Testing whether these patterns hold across multiple cities provides an opportunity to assess the extent of parallel intraspecific responses and identify macroecological drivers moderating these effects. We therefore investigated spider size, color traits, and web characteristics using urban-rural transects in 22 urban areas spanning the entire distribution range of *A. diadematus* in Europe and North-America. This involved recording in-field web trait measurements and capturing spider images to extract body size and color information through picture analysis. Using a meta-analytic approach, we investigated the response of these recorded phenotypic traits to urbanization, and related these responses to relevant city characteristics and potential environmental moderators.

## Methods

### Study design

From 2019 to 2022, 33 independent research groups searched for female spiders along urban-rural gradients in a total of 35 cities. Of these, 22 (17 European and 5 North American) were retained, while the others were excluded because no or insufficient spiders were found during searches (**Figure 1**; **Supplementary Table S1.1**). Urban areas were defined using the GHS-SMOD layer project (release 2023A, version 2.0, (Schiavina, Melchiorri, & Pesaresi, 2023) from the Global Human Settlement Layer (European Commission: Joint Research Centre, 2023). This layer contains footprints of Urban Centers (larger cities and metropolises) and Dense Urban Clusters (smaller cities and towns) drawn at 1000 m resolution using a methodology based on population density and built-up criteria (European Commission, 2021). Based on this, the 22 focal urban areas could be categorized in 18 Urban Centers and 4 Dense Urban Clusters, spanning a large portion of the current distribution range of *A. diadematus* (**Figure 1**). Among the 22 focal urban areas, 15 cities were surveyed for one year, while five areas were surveyed for two years and two areas for three years.

**Figure 1.**
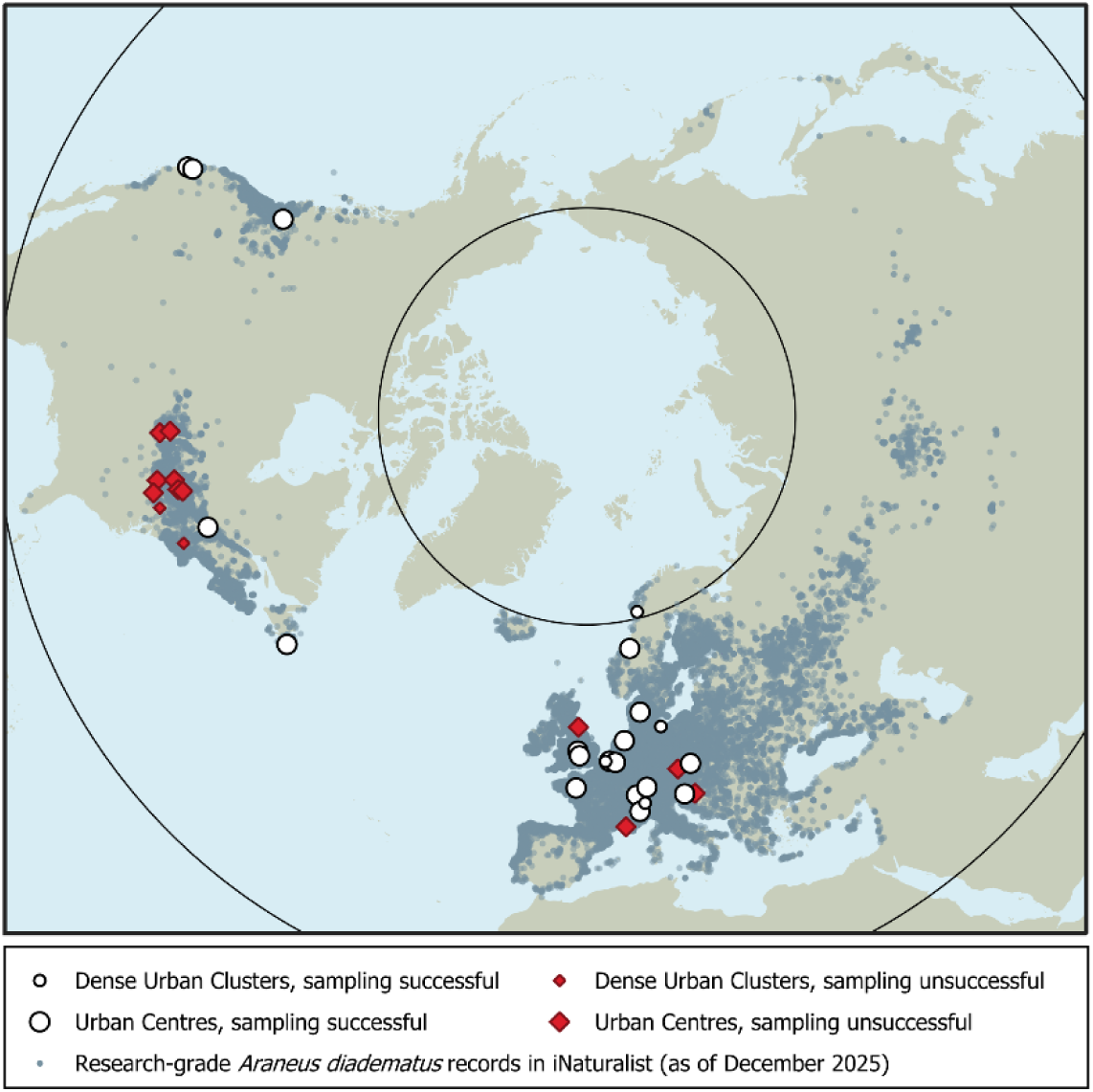
Map of the visited urban areas within the distribution range of *A. diadematus* (based on iNaturalist observations in GBIF; iNaturalist contributors & iNaturalist, 2025).

For the purpose of effect size calculations of urbanization on spider and web traits (see below), each unique combination of urban area and collection year was treated as a separate study, except the Vancouver (Canada) area that was visited in 2022 by two independent research groups. These two transects were in two distinct areas of the city, therefore, we treated these as two separate “studies” in our analyses.

### Spider and web trait collection

Spider and web traits were collected in sampling sites along urban-rural transects (see Supplementary Material 1 for more details). Web traits were measured directly in the field and spider size and color traits were measured digitally from standardized pictures taken in the field. Some spiders were not associated with an intact web, and for these spiders web trait measurements were missing. Pictures were taken by placing the spider in a Petri dish with an open lid, containing a measurement card. This card has a ruler on the edges and three gray scales (one large, two small) which were used during the image analysis for pixel and brightness calibration (see Supplementary Material 1). During analyses, pictures were inspected for correct spider orientation and lighting conditions, in the case of angled pictures in which the view of the entire abdomen was obscured or when the gray scale or spider were over- or underexposed, the affected size and/or color traits were not measured. Using this approach, measurements of up to five phenotypic traits were obtained per spider: body length, abdomen area (potentially related to fecundity or feeding status), abdomen brightness, web radius and number of crossing threads (**Figure 2**). We measured the web radius as the length of the upper part of the capture spiral, excluding the central hub and the potential area that is present above the capture spiral (see **Figure 2c**). An estimate of the average mesh size was obtained by dividing web radius by the number of crossing threads (**Figure 2d**). Measurements and pictures were submitted using the SpiderSpotter app, a citizen science app developed by Ghent University to register spiders and collect pictures (https://www.spinnenspotter.be/nl/app).

**Figure 2.**
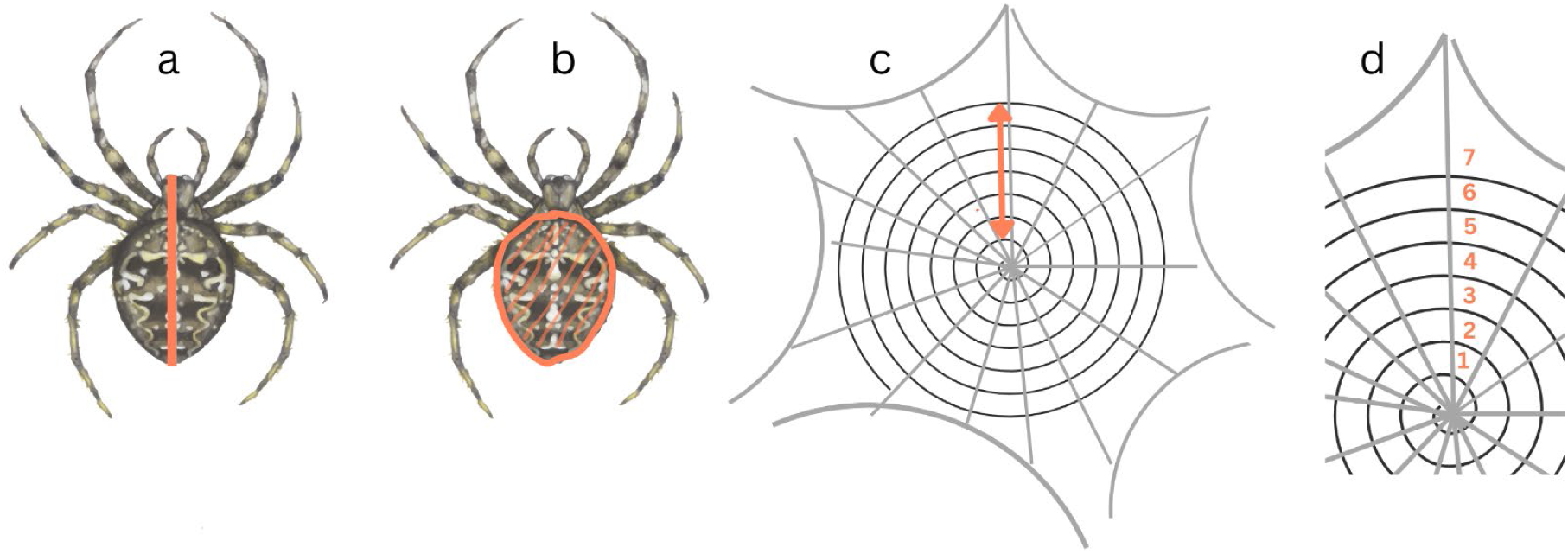
Graphical representation of the measured phenotypic traits: spider body length (a), abdomen surface area (b, this area was also used to measure abdomen brightness), web radius (c) and number of crossing threads (d); the latter two were used to calculate average mesh size.

From the 22 urban areas initially deemed sufficiently sampled, we obtained a total of 2110 raw spider presence records. We excluded one record due to geolocation errors that could not be corrected, one because it was collected in December after the main spider activity season, 19 because they were taken too far from their purported focal urban area (up to several hundreds of km), and 24 because they were not associated with trait data for any of the five traits of interest. This resulted in up to 2058 records potentially usable for analyses in 32 “studies”. As not all traits were collected or extractable from all records, the actual number of usable spider records and potential effect sizes varied between traits (**Figure 3**). To avoid biases, the compilation of the raw record database, including decisions on which photographs could be used for reflectance calculations, was undertaken by a different researcher (BV) than the one who set and applied criteria to keep or remove a record/study for subsequent analyses (MD).

**Figure 3.**
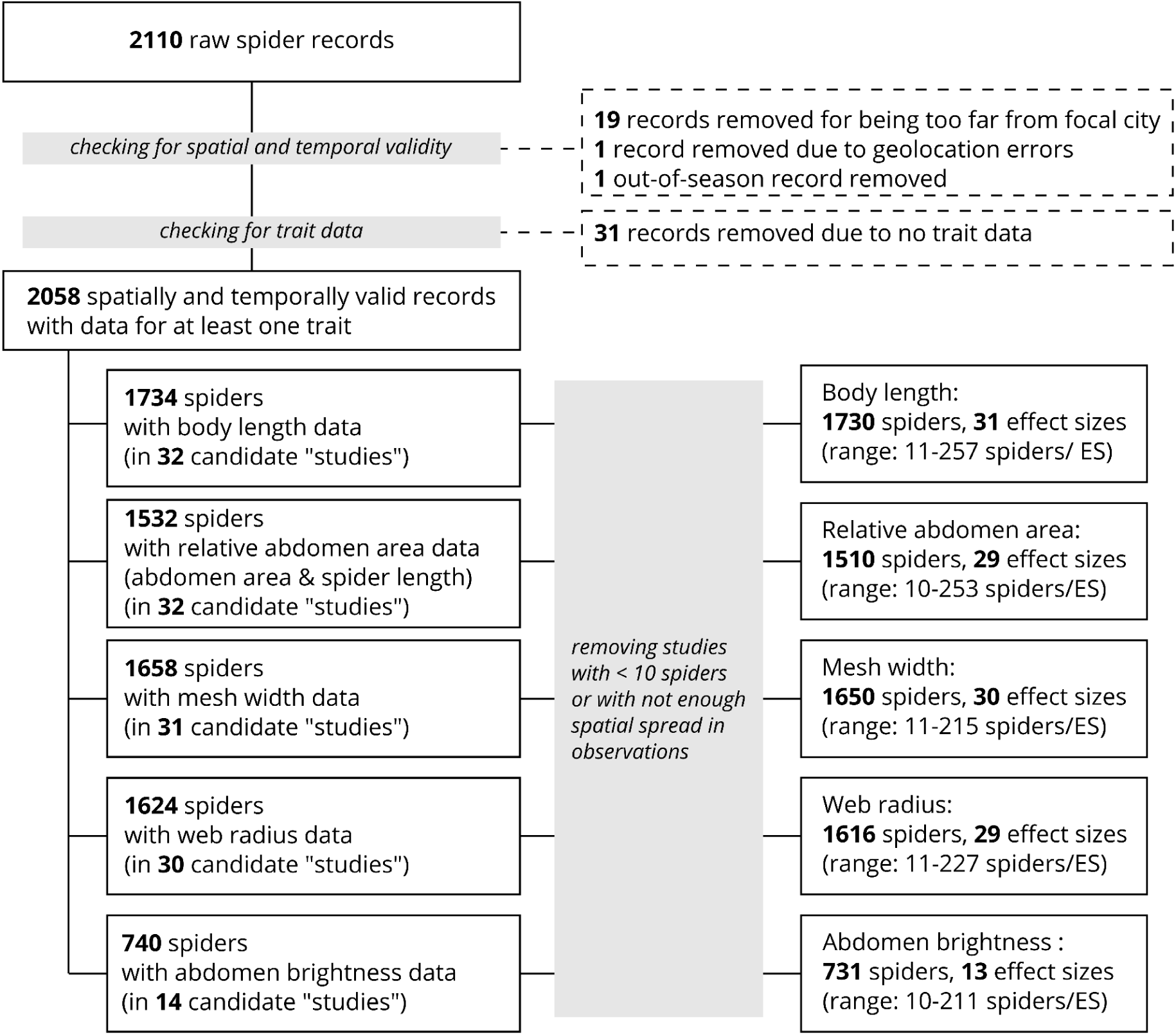
Flowchart summarizing data processing steps from raw records to effect sizes.

### Spider-level urbanization metrics

We used the Global Human Settlement Layer project to obtain urbanization data (European Commission: Joint Research Centre, 2023). We used the proportion of land surface covered by buildings (i.e. not counting roads and other artificial impervious elements) as our measure of urbanization intensity, taking built-up data from the GHS-BUILT-S raster layer (release 2023A), and the fraction covered by land from the GHS-LAND layer (release 2022A), both at the highest 10 m pixel resolution, which is available only for the reference year 2018, (Pesaresi & Politis, 2022, 2023). For each spider, we estimated the average proportion of built-up land in a range of circular buffers around its recorded location (50 m radius, and then 100 to 2000 m radius by 100 m increments). We estimated built-up proportions explicitly relative to land area rather than total buffer area because our dataset comprises several coastal cities. Radii wider than 2000 m were avoided *a priori* because, for some smaller cities and towns, such buffers would encompass the entire urbanization gradient, erasing differences between urban and non-urban sites. In addition, previous studies suggest the optimal scale of effect for spider responses is often under 2000 m (e.g. Bonte, Lanckacker, Wiersma, & Lens, 2008; Miyashita, Chishiki, & Takagi, 2012; Schmidt, Thies, Nentwig, & Tscharntke, 2008). Data extractions were done using the *sf* and *stars* R packages (Pebesma, 2018; Pebesma & Bivand, 2023).

### Moderators

In addition to spider-level urbanization metrics used to estimate effect sizes, we extracted several city-level variables (urban area size, mean annual temperature, mean Human Modification index, tree cover, mean sampling date) with the potential to explain differences between urban areas.

Many characteristics of cities relevant to their ecological functioning scale with city size (Norton, Evans, & Warren, 2016; Uchida et al., 2021), whether measured directly or through city population size, the latter often used in urban studies. This includes the magnitude of the urban heat island effect (Manoli et al., 2019), or street density and other characteristics of the street network (Levinson, 2012) which may drive habitat fragmentation. Furthermore, we may expect populations in larger urban areas to receive fewer immigrants dispersing from outside the city due to this fragmentation and distance, shaping gene flow and local adaptation (Richardson et al., 2014). We used the GHS-SMOD layer (see above) to delineate urban areas. For each urban area sampled, we collected the area of the focal GHS-SMOD polygon, subtracting areas covered by water based on GHS-LAND, as a measure of city size. We used log_10_-transformed values as a moderator (Manoli *et al*. 2019).

The responses of ectotherms such as spiders and their prey to urbanization could be dependent on regional climate (although evidence syntheses on abundance and richness are mixed, e.g. Fenoglio et al., 2020; Liang, He, Theodorou, & Yang, 2023; Vaz et al., 2023). Taking the centroids of the GHS-SMOD areas as reference points, we extracted mean annual temperature over the 1991-2020 30-year climatological period from the CRU-TS dataset, which interpolates data from weather stations into a 0.5° resolution grid (version 4.08, Harris, Osborn, Jones, & Lister, 2020). While post-2020 data are available from CRU-TS, there is at the time of writing, a slight decline in quality (number of stations available for interpolation) for one of the focal cities (St. John’s, Canada) after 2015 but especially after 2020. Precipitation may be another climatic driver of ectotherm responses, in addition to temperature. However, mean annual precipitation and mean annual temperature were significantly correlated across our sample of cities (with warmer cities being drier, **Supplementary Table S2.1**). Given this and given the relatively limited number of cities, we chose to use only temperature as climate moderator.

The effect of urbanization can also depend on the surrounding environment (Spotswood et al., 2021). For instance, if the environment around a focal city is already subject to heavy anthropogenic modification (both urban and non-urban, e.g. crops), there may be less contrast in suitability between that city and non-urban environments, shaping the observed impact of urbanization. As a synthetic indicator of the level of human modification of terrestrial habitats outside each focal city, we used the Global Human Modification dataset (Kennedy, Oakleaf, Theobald, Baruch-Mordo, & Kiesecker, 2019), which summarises data about multiple human impacts into a single continuous 0 (no human impact) to 1 (high human impact) metric (reference year 2016, 1 km resolution). We calculated the mean Global Human Modification value in a polygon made by drawing a 50 km radius buffer around the centroid of the focal GHS-SMOD area and then removing the area of the GHS-SMOD polygon itself from it.

Cities can vary widely in their greenness levels (Corbane et al., 2020; Volin et al., 2020), which can influence arthropod abundance and diversity (e.g. Turrini & Knop, 2015; Valdés-Correcher et al., 2022). Tree- and shrub-covered areas, and especially their edges, are favourable to *A. diadematus* (Hänggi, Stöckli, & Nentwig, 1995). As a measure of city greenness, we used the very high-resolution global tree cover data from (Tolan et al., 2024) and calculated the average proportion of land covered by trees within the perimeter of each GHS-SMOD urban area. Tree cover data were accessed and summarised using Google Earth Engine (Gorelick et al., 2017) and QGIS (QGIS.org, 2024).

Urbanization might have impacts on phenological dynamics, including phenological mismatches between interacting species (Fisogni et al., 2020; Theodorou, 2022). The effect of urbanization on predation-related traits (web characteristics) or body condition may thus depend on the time of the year. For the subset of spiders used in the estimation of each effect size, we collected the mean date of sampling, expressed in proportion of the year elapsed.

Across the 22 cities analysed, the five moderators we considered were for the most part not significantly correlated with each other. The only exception was a correlation between city size and average sampling date, with smaller cities being sampled earlier in the year (*r* = 0.59, **Supplementary Table S2.1**)

### Statistical analyses

All data analyses were done using R, version 4.4.2 and later (R Core Team, 2024). In addition to the specific packages cited elsewhere in this paper, we mainly used the *tidyverse* suite of packages for general data handling, cleaning and plotting (Wickham et al., 2019). We analysed data using a meta-analytic approach, treating input data from each city × year as its own separate “study” from which to extract effect sizes (as in e.g. Cote et al. 2022). Given that we are here specifically interested in the effects of urbanization, this approach avoids issues that would emerge by instead combining raw data from all cities in a single-step multilevel model (“mega-analysis”, e.g. Thompson et al. 2025). For instance, such a single-step model would require all cities to share the same urbanization predictor variable (i.e. same buffer width); it would not be possible to analyze responses to urbanization at the spatial scale at which they are the strongest/most relevant in each city (Quesnelle, Lindsay, & Fahrig, 2015).

### Computing effect sizes

For each study, we fitted a separate linear model to estimate the effect of urbanization (built-up proportion within a buffer) on each of the traits of interest. Not all traits were collected for all spider records, as e.g. in some cases spiders could be found without an intact web, or vice-versa. In addition, in some prospecting sessions/studies, not enough spiders were found along urbanization gradients to fit meaningful effect sizes, or they were only found in a narrow part of the prospected gradient, reducing the observed variation in urbanization. For each study × trait combination, we only fitted a model if there were at least 10 spider observations without missing data for the trait, and if these observations were sufficiently spaced from each other. To determine the latter criterion, we used the grid underlying the GHSL data at 100 m resolution, and considered observations sufficiently spaced out if they were spread over at least five 100 × 100 m pixels. Depending on the trait, this filtering removed between 1 (out of 14, abdomen brightness) and 3 (out of 32, abdomen area) potential studies from consideration (see **Figure 3**).

To test the effect of urbanization on body condition, we modelled log-transformed abdomen area as a function of both built-up proportions and log-transformed spider length, so the urbanization coefficients reflected the effect of built-up proportions after controlling for the expected allometric relationship between abdomen area and spider length, allowing urbanization effects to be evaluated independently of body size scaling. For all other traits we used simple univariate models linking the trait to built-up proportions. All models were fitted on centered and scaled data (Schielzeth, 2010) (scaling applied after log-transformation for the abdomen area models).

We fitted these linear models for each buffer radius at which urbanization was measured, and used AICc to select the best scale of effect for each study × trait combination (Quesnelle, Lindsay, & Fahrig, 2015; **Supplementary Figure S3.1**). For each of these selected models, we used the R package *DHARMa* (Hartig, 2022) to check whether model assumptions of residual spatial independence were met. If residual spatial autocorrelation was detected in a given model, it was refitted as a spatially explicit mixed model using the *spaMM* package (Rousset & Ferdy, 2014) to include a spatial random effect (using individual spider coordinates and a Matérn correlation structure).

Finally, we used the *t* statistics of the urbanization coefficients and degrees of freedom extracted from each retained model to estimate *Z*-transformed (partial) correlation coefficients *Z* as our effect sizes as well as their corresponding sampling variances *v̅* via the R package *metafor* (Aloe, 2015; Aloe & Thompson, 2013; Viechtbauer, 2010).

### Meta-analytic models

As effect size heterogeneity is expected in ecological data (Senior et al., 2016), we fitted random- and mixed-effect meta-analytic models (Koricheva, Gurevitch, & Mengersen, 2013; Nakagawa, Yang, Macartney, Spake, & Lagisz, 2023). Since multiple cities were visited several times, we further used a multilevel approach, with additional random effects of city identity, to account for this potential source of non-independence (Nakagawa et al., 2023). We fitted our models in a Bayesian context using the *brms* R package (Bürkner, 2017) as a frontend to the Stan language (Stan Development Team, 2024). Bayesian models make it easy to accurately propagate uncertainty in quantities calculated from model parameters (e.g. effect size heterogeneity and its partitioning). In addition, Bayesian meta-analyses tend to have better coverage properties and to be more conservative compared to frequentist equivalents when the number of effect sizes is low (Pappalardo et al., 2020).

For each trait, we first fitted intercept-only random-effect models, to estimate the overall meta-analytic mean effects. We further extracted the total random effect variances (between-cities variance + within-cities, between-studies variance = *τ*^2^ + *σ*^2^) and used them, along with effect size sampling variances *v̅*, to calculate the relative heterogeneities 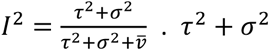 and *I*^2^ provide absolute and relative measures, respectively, of the spread of individual effect sizes around the average value (e.g. Nakagawa et al., 2023). Our hierarchical model also allowed us to estimate the proportion of total random effect variance linked to between-cities differences: *τ*^2^/(*τ*^2^ + *σ*^2^).

We then ran exploratory meta-regressions, adding to the above meta-analyses the variables described in **Moderators** above as fixed effects. Moderators were centered and scaled before inclusion (Schielzeth, 2010). We ran both multiple regressions including all five moderators at once (without interactions), and univariate regressions including one moderator at a time. We only ran the latter for abdomen reflectance, given the lower number of effect sizes available for that trait (**Figure 3**).

Priors were chosen to be weakly informative (*sensu* e.g. McElreath, 2020), with their scale adjusted to the plausible range of values for *Z*-transformed correlation coefficients and expected heterogeneities for ecological datasets (based on Senior et al., 2016; Röver et al., 2021). We used Normal(*μ* = 0, *σ* = 0.5) priors for fixed effect intercepts and moderator coefficients, as well as Half − Normal(0,0.4) priors for random effect standard deviations (see **Supplementary Material S4** and **Data and code availability** for an overview of the rationale; preliminary tests showed that results were not sensitive to slight changes in prior scales). We ran four chains per model, with 4000 iterations per chain and the first 2000 being used as warmup. We confirmed that bulk and tail effective sample sizes, as well as *R̂* values for checking convergence, were all satisfactory following Vehtari et al. (2021).

Whenever posterior summaries are provided, they are given as posterior means plus 95% credible intervals, using quantile intervals.

## Results

### General urbanization effects: intercept-only meta-analyses

Spiders in more built-up areas had consistently smaller abdomens relative to their body length across the urban areas studied (**Table 1**, **Figure 4**). The average effect size for the other four traits was not different from 0 (**Table 1**, **Figure 4**).

**Figure 4.**
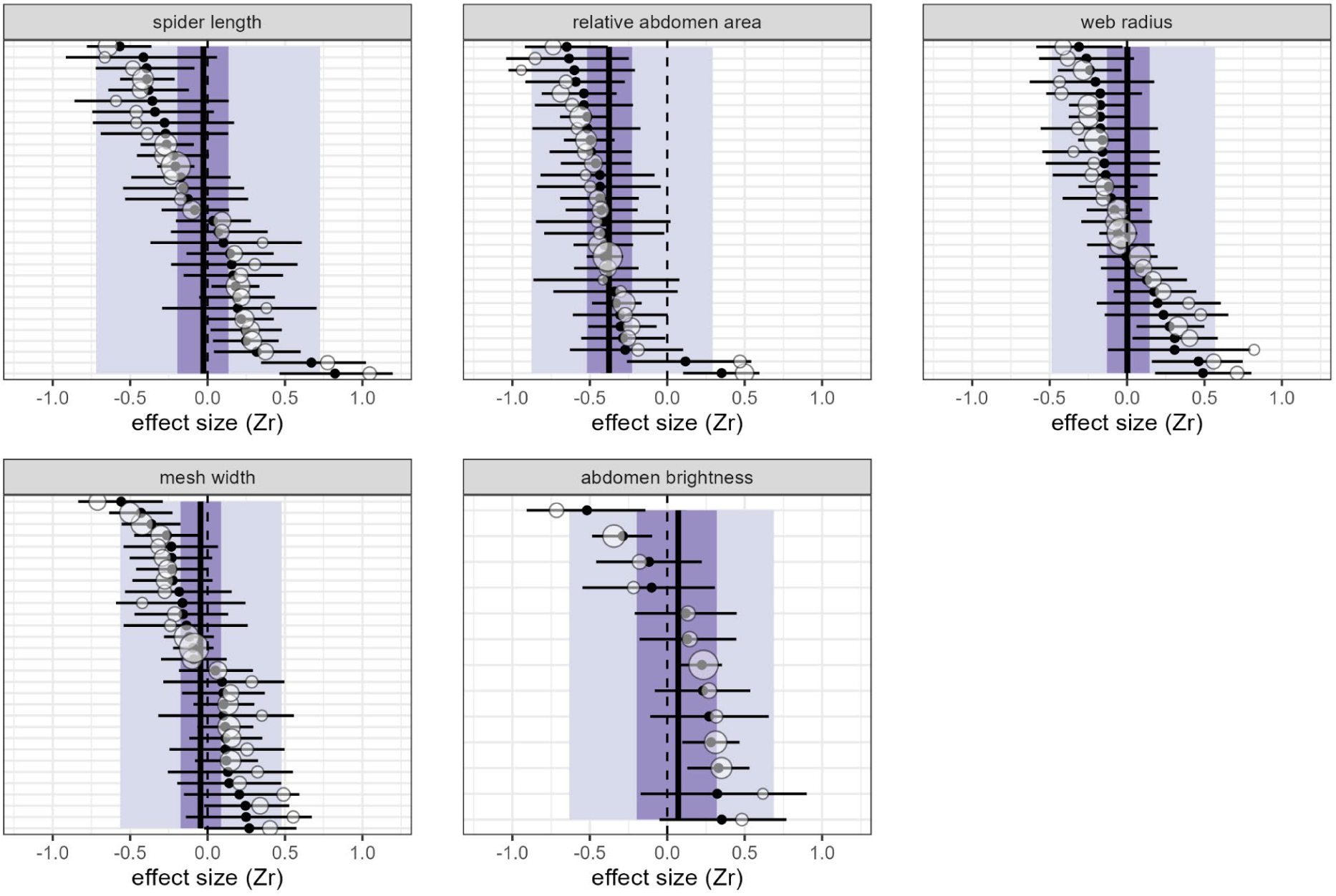
Observed and predicted effect sizes for all five studied traits. Effect sizes *Z* are ranked by increasing order of predicted values. Values > 0 indicate higher trait values when urbanization (built-up proportion) increases, values < 0 indicate lower trait values. White bubbles depict observed effect sizes with their width inversely proportional to sampling variance, while black dots and segments are predicted effect sizes and their 95% credible intervals from the models. The darker colored bands are the 95% credible intervals for the overall meta-analytic means, while the lighter colored bands are 95% prediction intervals, i.e. indicate the expected distribution of individual effect sizes. The large vertical black line depicts the predicted overall mean. Plots annotated with urban area and year information for each individual effect size are shown in **Supplementary Figures S5.3-S5.7**.

**Table 1.** Summary of the intercept-only random-effect meta-analyses. Notation for variance components follows Nakagawa et al. (2023).

|  | SPIDER LENGTH | RELATIVE ABDOMEN<br>AREA | MESH WIDTH | WEB RADIUS | ABDOMEN<br>BRIGHTNESS |
| --- | --- | --- | --- | --- | --- |
| predicted overall mean $Z_r$ | -0.03 [-0.19, 0.14] | **-0.38 [-0.52, -0.23]** | -0.04 [-0.17, 0.09] | 0.00 [-0.13, 0.15] | 0.07 [-0.20, 0.32] |
| absolute heterogeneity $\tau^2 + \sigma^2$ | 0.15 [0.08, 0.29] | 0.09 [0.03, 0.19] | 0.08 [0.03, 0.18] | 0.08 [0.04, 0.16] | 0.15 [0.04, 0.40] |
| relative heterogeneity $I^2$ | 0.87 [0.79, 0.94] | 0.78 [0.61, 0.90] | 0.79 [0.62, 0.90] | 0.79 [0.64, 0.89] | 0.85 [0.67, 0.95] |
| proportion of absolute heterogeneity associated with between-cities differences | 0.45 [0.00, 0.86] | 0.62 [0.01, 1.00] | 0.71 [0.03, 1.00] | 0.26 [0.00, 0.78] | 0.34 [0.00, 0.87] |

There was substantial trait effect size heterogeneity; posterior means for *I*^2^were > 0.75 across all traits (**Table 1**, **Supplementary Figure S5.1**). Accordingly, even in traits where the overall effect was not different from zero, predicted credible intervals for at least some individual effect sizes were frequently non-zero in either positive or negative direction (**Figure 4**). Trait variation along urban-rural gradients is thus strongly city-specific.

Attempting to partition between-studies variance into its between- and within-city components showed that for all five traits, posterior uncertainty is too high to determine reliably which component accounts for most of the heterogeneity (**Table 1**, **Supplementary Figure S5.2**).

### Macro-ecological drivers of variation in urbanization effects across the species’ range: meta-regressions

Using univariate meta-regressions, variation in effect sizes could be connected to a moderator in three of the five traits studied (**Table 2**). Urbanization affected web radius differently depending on the temperature (mean annual temperature): we observed larger webs in cities in colder regions, and a tendency for smaller webs in cities in warmer regions (**Figure 5**). Responses of two traits varied depending on the sampling date (**Table 2**). The effect of urbanization on relative abdomen size was more negative (smaller abdomens) when sampling occurred on a later sampling date (**Figure 6**), while the effect on abdomen brightness was more positive (paler abdomen colour) (**Figure 7**). Results from multiple regressions were qualitatively and quantitatively similar, with one exception: when other moderators were controlled for, mean sampling date was positively correlated with spider body length responses (**Supplementary Table S5.1**). Urbanization reduced spider body length when sampled on an earlier date, but this effect disappeared with later sampling dates (**Supplementary Figure S5.8**).

**Figure 5.**
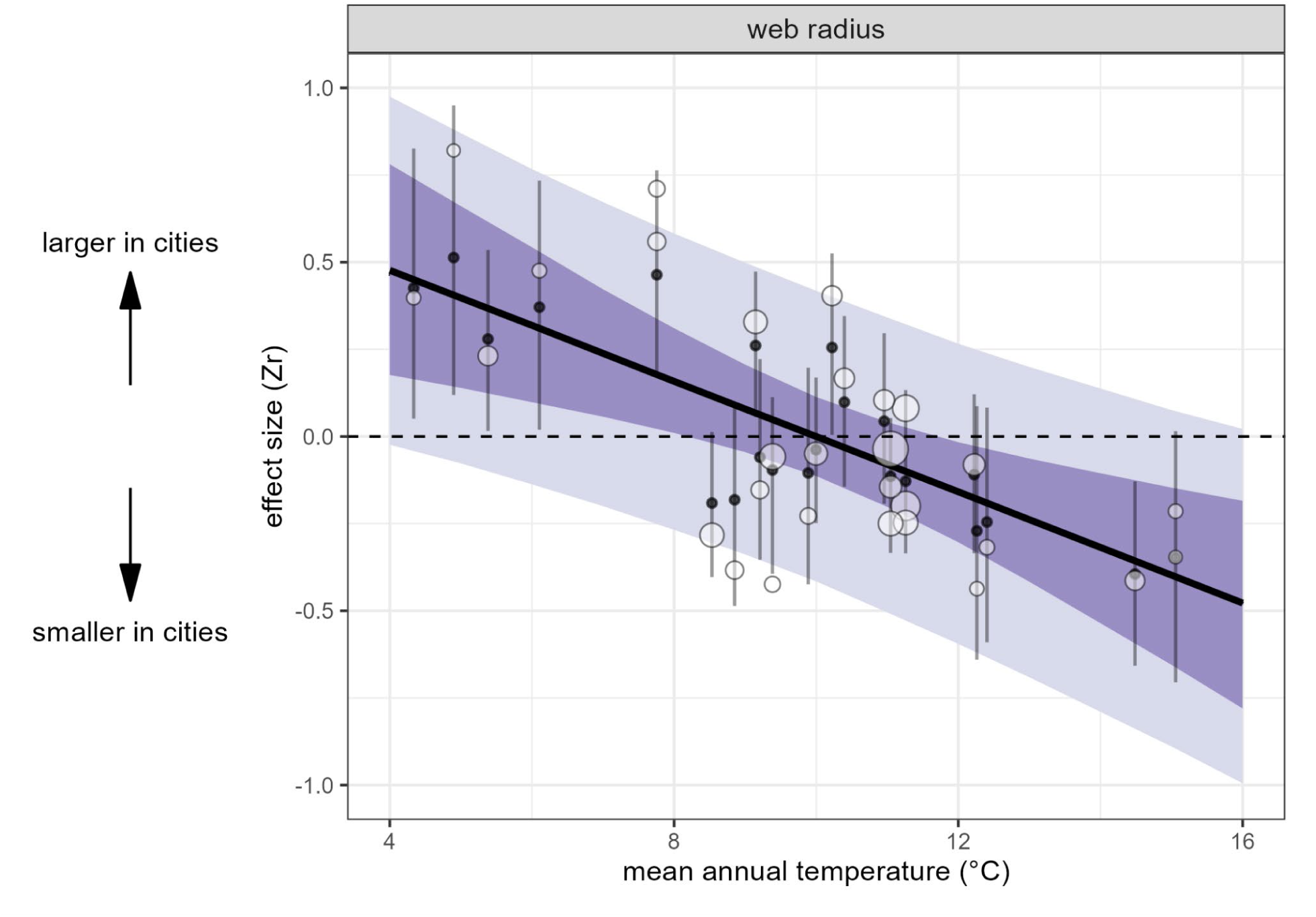
Effect of annual temperature on web radius response to urbanization. White bubbles are observed effect sizes with their width inversely proportional to sampling variance, while black dots and segments are predicted effect sizes and their 95% credible intervals from the model. The darker colored band represents the 95% credible interval for the overall meta-analytic mean regression line, while the lighter colored band denotes 95% prediction intervals, i.e. indicates the expected distribution of individual effect sizes.

**Figure 6.**
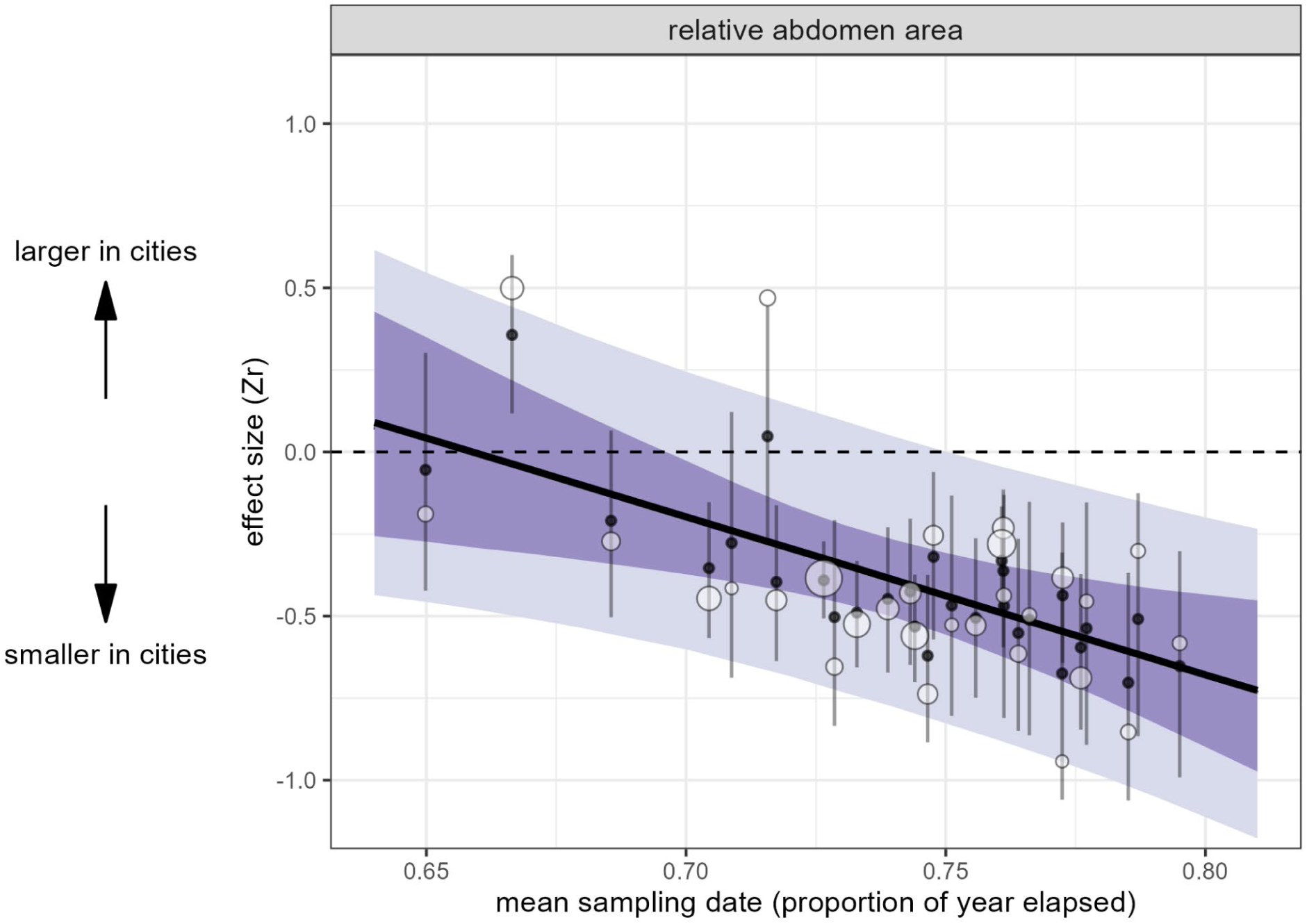
Effect of sampling date on relative abdomen area response to urbanization. Average sampling dates in studies with effect sizes range from 25th of August (2021) to 17th of October (2022). White bubbles are observed effect sizes with their width inversely proportional to sampling variance, while black dots and segments are predicted effect sizes and their 95% credible intervals from the model. The darker colored band represents the 95% credible interval for the overall meta-analytic mean regression line, while the lighter colored band denotes 95% prediction intervals, i.e. indicates the expected distribution of individual effect sizes.

**Figure 7.**
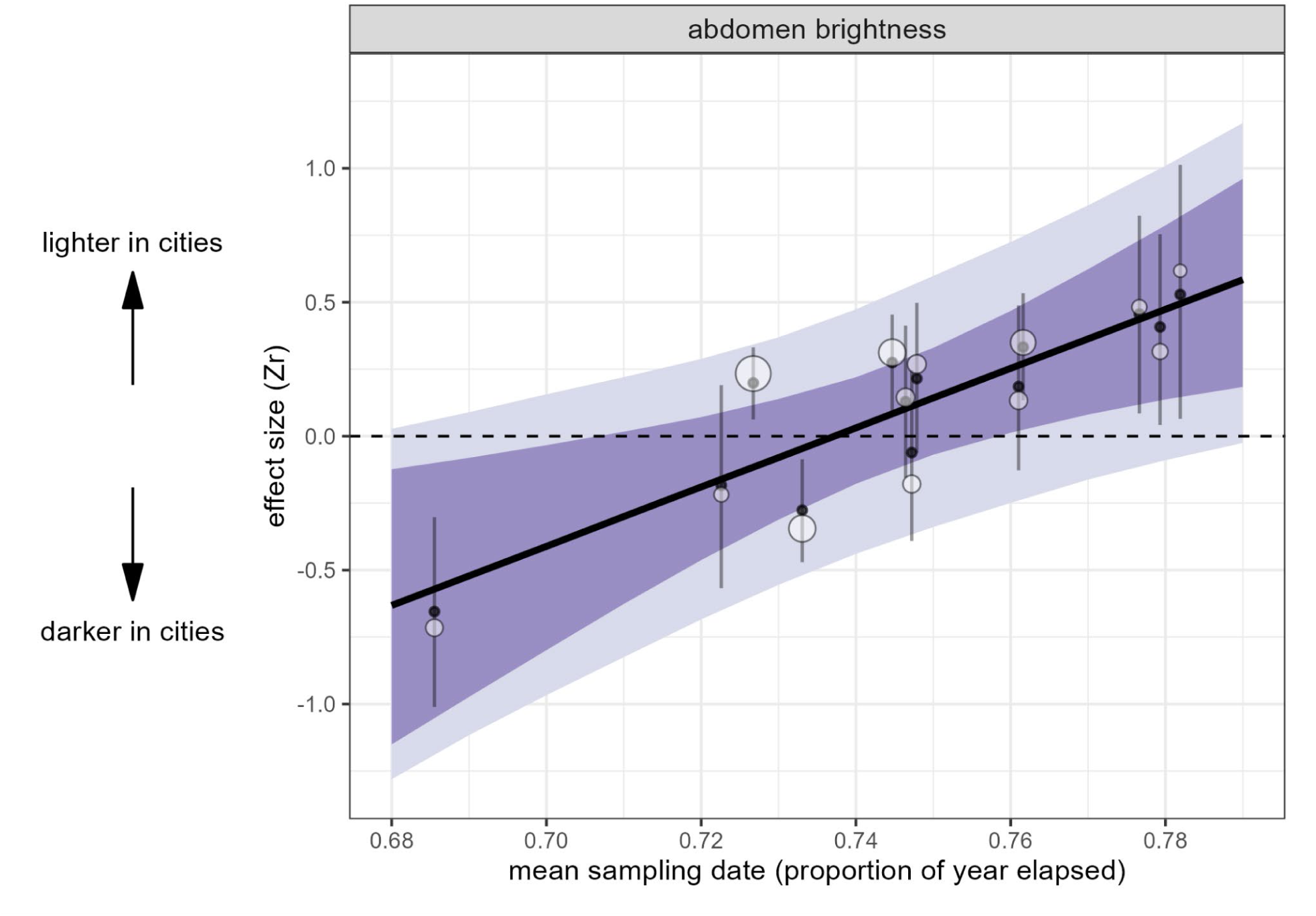
Effect of sampling date on abdomen brightness response to urbanization. Average sampling dates in studies with effect sizes range from 7th of September (2022) to 12th of October (2022). White bubbles are observed effect sizes with their width inversely proportional to sampling variance, while black dots and segments are predicted effect sizes and their 95% credible intervals from the model. The darker colored band represents the 95% credible interval for the overall meta-analytic mean regression line, while the lighter colored band denotes the 95% prediction interval, i.e. indicates the expected distribution of individual effect sizes. Note that the range of dates on the x-axis is narrower here than in Figure 6, due to fewer valid effect sizes for this trait.

**Table 2.**
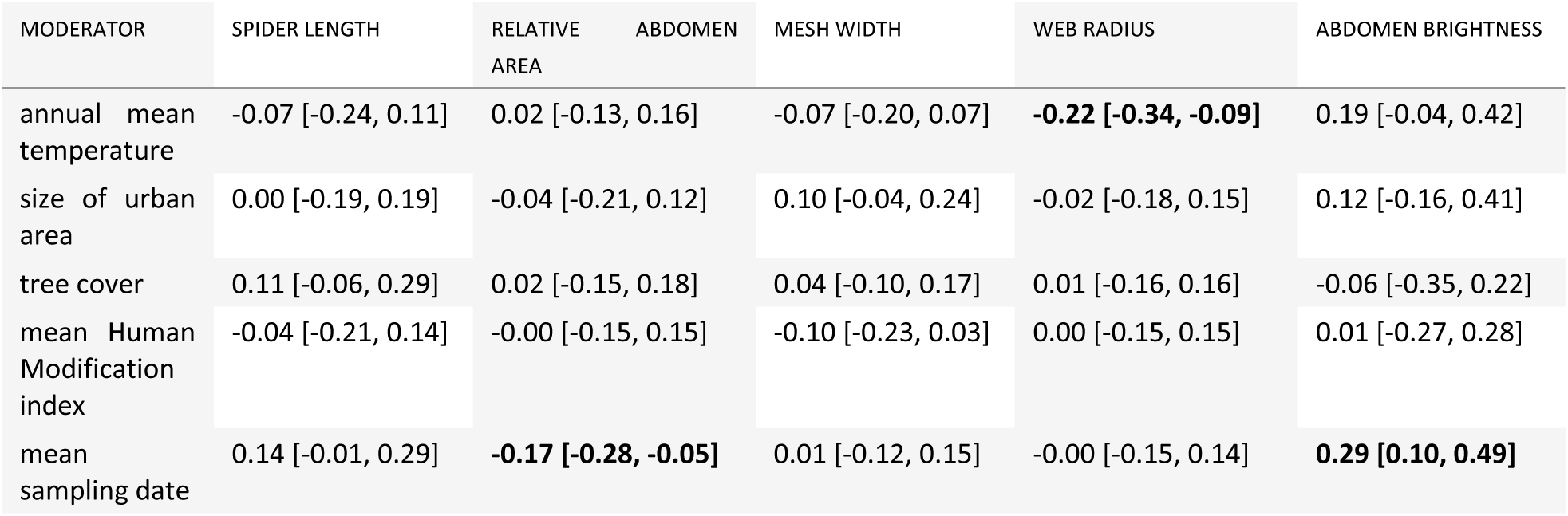
Summary of moderator effects from meta-regressions. Moderators, but not effect sizes, were centered and scaled to unit 1 standard deviation before model fitting. Coefficients are taken from univariate models including one moderator at a time (see **Methods**); results from multiple regressions are qualitatively and quantitatively similar and presented in **Supplementary Table S5.1**.

## Discussion

Urbanization often homogenizes ecological conditions, making cities more similar to each other than to surrounding non-urban areas (Groffman et al., 2014). This has led to the assumption that cities exert comparable selective pressures and that species responses should be uniform (a “cities as replicated natural experiments” perspective). However, multi-country studies with replicated sampling (Cosentino & Gibbs, 2022; Santangelo et al., 2022) reveal that urban trait responses are strongly contingent on the prevailing environmental context and climate. Because phenotypes reflect both ecological and evolutionary processes, variation in urban responses is expected and shaped by local conditions influencing morphology, physiology, life history, and behavior. Our meta-analysis confirms this complexity: spider trait responses to urbanization were highly city-specific, with most traits showing no consistent pattern across cities. For every trait, at least one city exhibited a clear effect in either direction. Responses were also trait-specific. Urban living consistently reduced abdomen relative surface area, an effect amplified when spiders were sampled later in the year. Only web-building showed a clear environmental moderator: the urbanization effect on web radius correlated strongly with mean annual temperature. For body length and abdomen brightness, investigated moderators did not explain inter-city variation, though body length of urban spiders was higher relative to non-urban when sampled on a later date. These findings underscore that urban evolutionary and ecological dynamics are context-dependent, highlighting the need to incorporate local and climatic factors when assessing species responses to urbanization.

Among the pressures associated with urbanization, increased temperatures, habitat loss, and altered species interactions can shift communities toward smaller-bodied species (but see Martin & Sheridan, 2022; Merckx et al., 2018). Similar size reductions may occur within species, though responses are taxon-specific (Cabon et al., 2024; Martin & Sheridan, 2022). In *A. diadematus*, body size reduction in urban populations was previously documented along urban–rural gradients in Flanders (Dahirel et al., 2019). That study used cephalothorax width as a proxy for size, whereas we applied a composite measure incorporating the length of both the cephalothorax and abdomen. While sclerotized structures remain constant in adults, abdomen size varies with feeding and reproductive state (Jakob et al., 1996). Accordingly, we observed a similar negative response of relative abdominal surface area in most cities. Because abdomen size correlates with fecundity (Higgins, 1992; Marshall & Gittleman, 1994; Wise, 1975), this suggests urbanization imposes strong negative effects on reproductive potential. These findings align with Dahirel et al. (2019), who reported fewer eggs in urban females, even though spider abundance remained comparable between urban and rural sites.

Spider body length showed highly variable responses across cities, with many non-zero effect sizes. Contrary to expectations of smaller size, increases were as common as decreases, indicating no consistent size shift. Larger body size in urban spiders has been reported elsewhere (Ripp et al., 2018), often linked to greater prey availability in cities (Lowe et al., 2016). Despite selecting moderators expected to influence size, none explained the observed heterogeneity among cities. Instead, we found an effect of sampling date on the relationship between spider length and urbanization, with more positive urbanization effects (longer urban spiders) observed later in the year.

Some caution needs to be taken when interpreting sampling date as indicative of early or late time periods in a year, as higher latitude affects spider phenology through shorter growing seasons that start later and end earlier. This results in *A. diadematus* taking two years to mature in northern regions (Toft 1976). We find a significant negative correlation between average sampling date and latitude (**Supplementary Table S2.1**) indicating that northern cities were sampled earlier.

Observing longer urban spiders later in the year can be explained by increased abdomen length due to egg production in urban spiders, by differential mortality with a relatively higher mortality rate of smaller and early maturing spiders, or by an increased proportion of larger and later maturing females (Šet *et al*. 2021) during fall in urban areas. However, our data do not support the first hypothesis as relative abdomen size decreased with urbanization later in the year and indicates that an increased overall size at maturity does not imply higher fecundity, likely because of a reduced prey availability or increased energetic investments to cope with urban conditions (e.g. reduced activity if increased temperatures reach thermal limits, desiccation,…).

Abdomen shape may be an additional important factor in interpreting these body size patterns. A longer but narrower abdomen would increase total spider length while simultaneously reducing abdomen area. Such an elongated abdomen could be selected in hot, urban environments, as it has been shown to reduce the occurrence of high body temperatures in spiders (Ferreira-Sousa, Rocha, Motta, & Gawryszewski, 2021). It is interesting to note that, for example, in Bern, Switzerland, spiders exhibited increased body length with increasing urbanization (**Supplementary Figure S5.3**), while exhibiting comparative reduction in abdomen surface area (corrected for body size), indicating that spiders might indeed become longer, but thinner in some cities.

We expected strong urbanization effects on spider body size due to reduced prey availability (Dahirel et al., 2019) and higher metabolic demands under urban heat island conditions (Oke, 1982). However, our results reveal a more complex, trait-specific pattern: only abdomen surface area consistently decreased, while body length showed heterogeneous responses. This aligns with previous work demonstrating taxon-dependent size responses and interactions with life history (Cabon et al., 2024), climate (Hantak et al., 2021), and latitude (Beasley et al., 2018; Lövei & Magura, 2022). The pronounced heterogeneity in body length across cities and years may reflect temporal variation in temperature or humidity, as well as local differences in microclimate, weather, and habitat structure (Cabon et al., 2024), all of which influence thermoregulation and prey availability.

Orb webs can be considered an extended phenotype, one that is often closely linked to body size, both because resources available for web building scale with size (Gregorič, Kuntner, & Blackledge, 2015) and because spiders may use their own body (front legs) as a yardstick when building (Vollrath, 1987). Given this link, we expected spider size and web traits to respond similarly to environmental changes. Previous research has shown that web size responds to land-use changes in ways that optimize foraging efficiency, though these responses can vary (smaller or larger webs in urban areas) (Bonte et al., 2008; Dahirel et al., 2019; Lowe et al., 2016; Ripp et al., 2018). For *A. diadematus*, urban spiders in Belgium built smaller, finer-meshed webs in response to local scale urbanization, whereas web area increased with landscape-scale urbanization, consistent with behavioral responses to smaller prey in urban environments (Dahirel *et al.,* 2019). In this study, we observed macrobehavioral patterns (large-scale behavioral variation across space and time; Keith, Drury, McGill, & Grether, 2023) in web-building, with city-dependent increases and decreases in web and mesh size in urban areas. Web sizes increased with urbanization in colder regions but decreased in warmer regions, a relationship that does not match any pattern in the observed responses of body size. In contrast, responses in mesh size could not be explained by the investigated moderators. If the increase in web size in cities in colder regions is a response to smaller prey, this suggests that urbanization effects may be stronger in colder regional climates compared to warmer ones, and could suggest a potentially more pronounced effect of the local urban heat island in cold cities on prey availability. A smaller prey body size in northern latitudes is consistent with latitude-related size patterns in insect communities (converse Bergmann rule; Blanckenhorn & Demont, 2004) and with evidence that climate and urbanization jointly modulate body size responses (Lövei and Magura, 2022; Hantak et al., 2021). For example, carabid beetles demonstrate smaller body sizes with higher latitudes, a pattern that is more pronounced in urban areas compared to rural ones (Lövei and Magura, 2022). Whether urbanization effects result in differing patterns of prey availability between northern and southern cities warrants further investigation. This is especially true as mesh size responses were not consistent across cities and could not be explained by the investigated moderators, despite previously having been shown to respond to local urbanization levels in Belgium for *A. diadematus* (Dahirel *et al*. 2019). This suggests that other factors (including microhabitat characteristics) might play a role in shaping web responses. Regardless of the precise mechanisms driving the observed macrobehavioral variation in web-building, these changes likely have top-down effects on local food webs (Keith et al., 2024). This further reinforces the idea that intraspecific variation in functional traits must be accounted for to understand community-level responses to urbanization (Des Roches et al., 2021; Des Roches et al., 2018; Raffard, Santoul, Cucherousset, & Blanchet, 2019).

Among all investigated traits, we expected consistent urban-rural shifts in spider coloration due to its demonstrated role in thermoregulation and camouflage, in coping with pollution and due to potential physiological constraints on pigment production linked to reduced resource availability in urban environments (Leveau, 2021; Stuart-Fox et al., 2017; Trullas et al., 2007). Indeed, among ectotherms, urban-rural shifts in color can align with thermoregulatory demands. *Cepaea nemoralis* snails tend to have lighter shells in cities which is consistent with the urban heat island effect as lighter individuals heat up less compared to darker counterparts (Kerstes, Breeschoten, Kalkman, & Schilthuizen, 2019). As with Dahirel *et al*. (2019) for spiders in Flanders, this study was limited to a single region (Netherlands). In our current study, despite substantial heterogeneity within and among locations, and our study spanning a large range of the climatic conditions experienced by *A. diadematus*, we did not observe a systematic effect of urbanization on spider color and no evidence that color responses varied with regional temperature. We found that spiders collected in cities later in the year were paler, which may reflect either selective survival of brighter individuals in urban areas or plasticity in coloration. In *A. diadematus* color is produced through two distinct mechanisms: white markings, including the characteristic cross pattern, are formed by guanine crystals (Seitz, 1972), while the light to dark background coloration is likely pigment-based. Although guanine crystals derive from nitrogen metabolism in spiders (Levy-Lior et al., 2010; Nentwig, 1987), we detected no clear link between moderators potentially associated with reduced resource availability along the urban gradient (tree cover, Human Modification Index) and color variation. Guanine crystals have been found to play a role in color change in spiders (Wunderlin & Kropf, 2012) with some evidence for color plasticity in *A. diadematus* (Blanke & Merklinger, 1983, but see Messas *et al.,* 2025). Recent evidence points towards a camouflage role of body coloration in *A. diadematus* by background matching (Messas *et al.,* 2025) which could explain the variable responses in our study as background coloration of urban and rural web locations wasn’t included as a potential driver.

Our study demonstrates that urbanization induces both morphological and behavioral responses in the common garden spider. While earlier research suggested more uniform responses across cities from a single region, our replicated surveys across the Northern Hemisphere and meta-analytical approach reveal substantial heterogeneity, even after accounting for key moderators. Of all traits, only relative abdomen surface area showed a consistently negative response across most cities, becoming more pronounced later in the season. Web size responses were moderated by climate: spiders built larger webs in colder cities and smaller webs in warmer ones.

These findings highlight that, despite global homogenization of urban environments (Piano et al. 2020; but see Lokatis & Jeschke, 2022), intraspecific responses remain largely context-dependent, shaped by climatic and phenological factors. Future work should integrate finer-scale urbanization metrics and local environmental data to disentangle drivers of this variability (Moll et al., 2019; Szulkin et al., 2020). Combining broader replication across cities with more intensive within-city sampling will help clarify general patterns and mechanisms underlying phenotypic responses to urbanization.

## Data availability

Data and R code needed to run all analyses presented in this study are available as a GitHub repository (https://github.com/mdahirel/SPINCITY-scientists-2024), which is archived in Zenodo (DOI: https://doi.org/10.5281/zenodo.14809653). Note that large raw files directly from external archives (e.g. GHSL, CRU-TS, see Methods for references) are not included, but links to these archives, code to process them and datasets produced from them are present in the repository.

## Acknowledgement - funding

This study was supported by the Research Foundation - Flanders (FWO, grant G080221N). We would like to thank Barbara Helm for her coordinating efforts in this project.

