## supplementary material for "Urbanization effects are trait-specific and city-dependent across a widespread spider’s global range"

### Supplementary Material 1: details on the transect design and image analysis for spider color

#### Transect design

In each city (Table S1.1) we aimed to sample spiders along a clearly delineated urban-rural transect, though local variations made this difficult sometimes; in these cases spiders were sampled more haphazardly along the general urban-rural gradient. The teams aimed to perform spider sampling in the shortest possible time frame while avoiding unfavorable conditions such as wind and rain, that hinder locating European garden spiders and might cause damage to webs.

Table S1.1. Overview table of the urban areas visited, with visit year denoted by "x" when sampled successfully and by "-" when visited but unsuccessfully sampled. All sampled area-year combinations were sampled by one transect, with the exception of Vancouver (*) in 2022 which was sampled by two independent research groups. We regard these two transects as two separate sampling events.

| Urban area | Country | 2020 | 2021 | 2022 |
|---|---|---|---|---|
| Gent | Belgium | x | x | x |
| Leuven | Belgium | | x | |
| Torhout | Belgium | x | x | x |
| Montreal | Canada | | x | |
| St. John's | Canada | | | x |
| Vancouver | Canada | | | x (*) |
| Brno | Czech Republic | | | x |
| Aarhus | Denmark | x | x | |
| Rennes | France | | x | |
| Greifswald | Germany | | | x |
| Konstanz | Germany | | | x |
| Torino | Italy | | | x |
| Verbania | Italy | | | x |
| Groningen | Netherlands | | x | |
| Bodø | Norway | | x | |
| Trondheim | Norway | | | x |
| Ljubljana | Slovenia | x | x | |
| Bern | Switzerland | | x | x |
| Oxford | UK | | x | |
| Warwick | UK | | | x |
| San Francisco Bay Area | USA | | x | x |

| | | | | |
|---|---|---|---|---|
| Santa Cruz | USA | | x | |
| Hamilton | Canada | | - | |
| London | Canada | | - | |
| Toronto | Canada | | - | |
| Zagreb | Croatia | | | - |
| České Budějovice | Czech Republic | | - | |
| Aix-en-Provence | France | | - | - |
| Newcastle upon Tyne | UK | | | |
| Akron | USA | | - | |
| Amherst (Massachusetts) | USA | | - | |
| Chicago | USA | | - | |
| Milwaukee | USA | | - | |
| Pittsburgh | USA | | - | |
| State College (Pennsylvania) | USA | | - | |

Transects were established prior to field sampling and were generally designed to start in the city center and extend towards a rural, green area outside of the city borders, but also considered accessibility of the locations. The tentative transect length was determined by measuring the distance from the city center to the city border and adding this same distance to the transect part in rural areas. This way, approximately half of sampling locations are located in the city and half outside, achieving a balanced sampling across urban and rural locations. Most cities were sampled only in one year. To determine the sampling locations across the transect, transect length was divided by 40, resulting in the distance between sampling locations, (for example, a transect of length 8400 meter would result in sampling locations every 210 meter). Starting from the beginning of the transect in the city center, these 41 sampling locations were then plotted on the transect. If the location was inaccessible, the nearest possible suitable location was selected for sampling. Upon arriving at a sampling site, a total of 20 minutes search time (divided over all team members) was allocated to search for spiders in a 100 meter radius. If no spiders were found, the next location was sampled. If spiders were present, web traits were first measured as catching the spider from the web can result in web damage. Web radius and the number of crossing threads were measured/counted. Web radius was defined as the length of the upper part of the capture spiral, without the central hub and the area above the capture spiral. Dividing web radius by the number of threads gave us the average mesh size.

The spider was then collected from the web and a picture was taken in a Petri dish with a custom-made Spiderspotter measuring card (see **Supplementary Figure 1.1**). This card contains a ruler on the edges and three gray scales (one large, two small) which were used during the image analysis. Card dimensions are 5.5 cm width and 8.5 cm length and consists of coated paper, 350 gr. Instructions advised taking pictures under a 90° angle to reduce variation in length measurements, and during suitable lighting conditions to avoid over- or underexposure and equal lighting of both the gray scale and spider. Using image analysis (see below), the following measures were taken: body length, abdomen surface area and abdomen brightness.

**Image analysis**

Image analysis was performed in ImageJ (version 1.54g.). During analyses, pictures were inspected for correct spider orientation and lighting conditions. If picture angle and lighting exposure was inappropriate, the corresponding affected size and/or color traits were not measured. This resulted in a different number of observations for each trait as, for example, certain pictures allowed for size but not for color measurements. Similarly, webs might be damaged in the field, not allowing for web measurements, while corresponding spider pictures could be taken. Pictures were calibrated by measuring one cm on the card using the "straight line" tool and setting the number of pixels contained by this line to 10 mm (Under "Set Scale"). The same "straight line" tool was used to measure spider length (**Figure 2a**). The "polygon selection" tool was then used to trace the edge of the abdomen to obtain abdomen surface area. This also enabled us to calculate the brightness of this area by averaging the values of the R, G and B channels ((R+G+B)/3). Next, one of the gray scales on the card was selected in such a way that lighting conditions maximally resembled those of the spider and with suitable exposure. To calculate the brightness of this gray scale, three points were placed into each of the gray scale squares using the "multipoint" tool. Per square, these brightness values were averaged and plotted against the reflectance of the gray squares, obtained from lab measurements (see below). This results in an exponential calibration curve which can be used to convert the measured abdomen brightness values into calibrated reflectance values (Johnsen, 2016).

We determined reflectance values for black, white and each of the gray rectangles using spectrophotometry. Diffuse reflectance was measured using a AvaSpec-2048 spectrometer and a dual light source set-up (AvaLight-DH-S deuterium halogen and AvaLight-HAL-S-MINI light source) with a bifurcated probe and integrating sphere (AvaSphere-50-REFL). A black reflectance standard (BS-2, Avantes) and white standard (WS-2, Avantes) was used for standardization. We measured three replicates of each rectangle per card, for five randomly selected cards in total. Reflectance spectra were processed using the pavo2 package in R (Maia et al. 2019) and average reflectance (%) for each rectangle was calculated. These values were further averaged over all five cards to obtain average reflectance values for all gray scales.

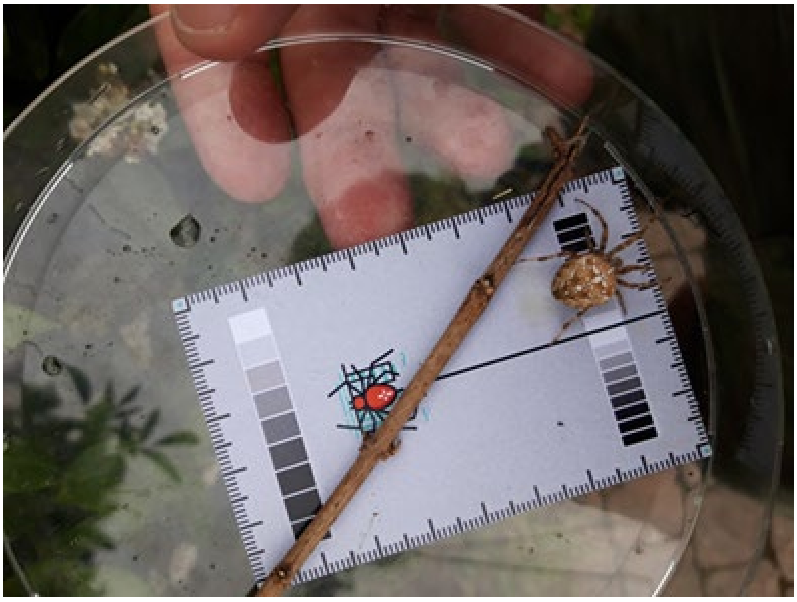

Figure S1.1. Example of a spider picture taken in the field, with the Spiderspotter card.

#### Supplementary Material 2: Correlation between potential moderators

**Table S2.1.** City-level (*N* = 22) Pearson correlation coefficients between the five *a priori* chosen moderators, plus city latitude and precipitation. For average sampling dates, the value used for cities visited multiple times is the mean of the different visits. Significant correlations are in bold; *: p <0.05, **:p<0.01; ***:p<0.001.

| | Temperature | $log_{10}$(city area) | Human Modification Index | Tree coverage | Average sampling date | Latitude |
|---|---|---|---|---|---|---|
| $log_{10}$(city area) | 0.15 | | | | | |
| Human Modification Index | 0.32 | -0.04 | | | | |
| Tree coverage | -0.04 | 0.32 | -0.12 | | | |
| Average sampling date | 0.32 | **0.59**** | 0.24 | 0.09 | | |
| Latitude | **-0.69***** | **-0.45*** | -0.09 | -0.29 | **-0.54**** | |
| Precipitation | **-0.52*** | -0.02 | **-0.64**** | 0.11 | -0.23 | 0.24 |

#### Supplementary Material 3: Scales of effect

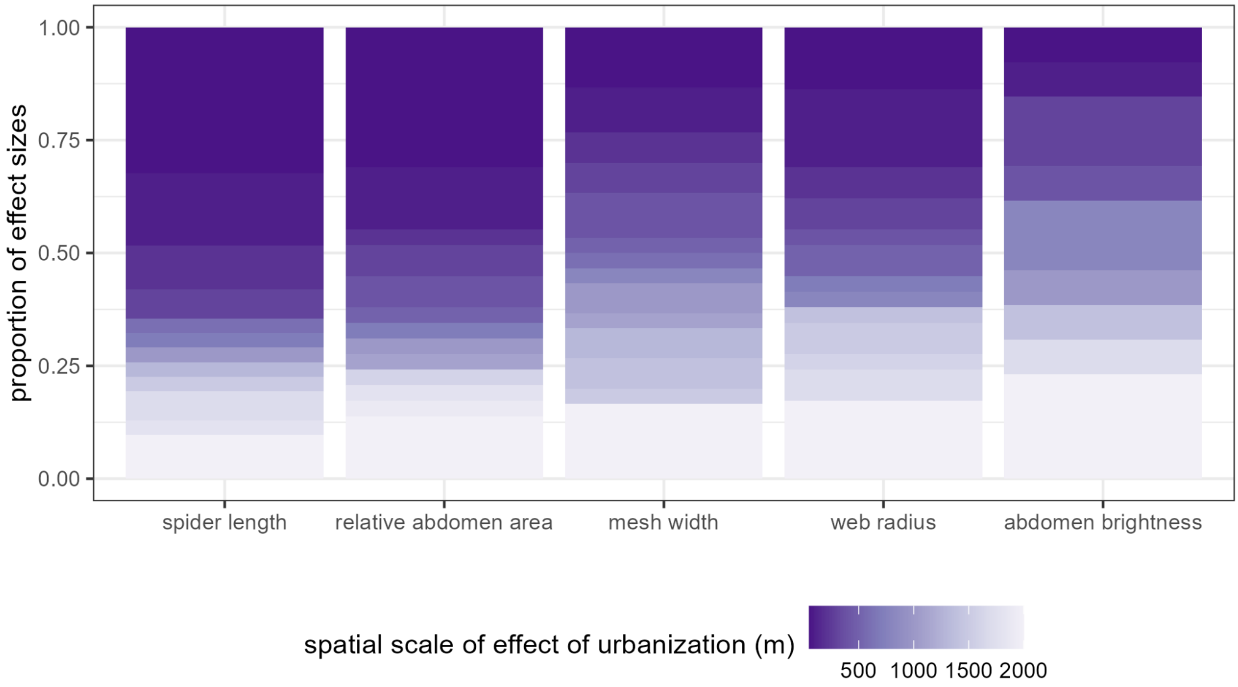


**Figure S3.1.** Distribution of the spatial scales retained for the effect of urbanization, by trait. Darker colors correspond to cities where the retained radius at which urbanization affects the spider trait is smaller (local scale), lighter colors cities where it is larger (closer to landscape scale).

### Supplementary Material 4: Prior predictive checks

As documented in Senior et al. (2016), the typical relative heterogeneity $I^2$ for ecological data is expected to be high, with the median estimate from their meta-analysis of meta-analyses being 84.67% and their mean estimate 91.69%. This directly gives us quantitative prior information with which to do prior predictive checks (McElreath et al. 2020).

We generated prior distributions of $I^2$ assuming various values of the scale parameter σ for the prior on random effect standard deviations. We assumed that the total random effect (between-studies) variance was split equally between the two random effect levels (among-cities and within-cities), with both levels' SD drawn from the same Half-Normal(0,σ) distributed prior, as mentioned in the main text. $I^2$ depends on both between-studies variance and the average study sampling variance $\bar{v}$; we used for these checks the mean $\bar{v}$ of all five traits (≈ 0.02; all traits had very similar values; **Data and code availability** in main text).

Based on this, we find that assuming a prior scale σ of 0.4 provides the most satisfactory matches to Senior et al. (2016). While a prior σ of 0.3 matches better the median estimate, it deviates more strongly from the mean estimate (**Supplementary Figure S4.1**, left). Larger σ values tend to overshoot both mean and median $I^2$, while smaller values lead to underestimations.

Regarding the prior variance of fixed effects, Röver et al. (2021) suggest that using total between-studies variance (i.e. the sum of among- and within-cities variance here) is a valid choice. Drawing on the simulations above, this results in a prior SD of about 0.5 for fixed effects, if the prior for random effect scale is set to 0.4 (**Supplementary Figure S4.1**, right).

While obtained independently through the use of ecology-specific pre-existing knowledge, such prior choices closely match some of the suggestions made by Röver et al. (2021) on more general grounds: they suggested that a prior between-study variance of 0.3 would be suitable for high heterogeneity cases, resulting in a total between-studies SD of ≈ 0.55.

We note that results presented in the main text are nonetheless robust to small changes in the prior (not shown, but can be tested using the code in **Data and code availability**).

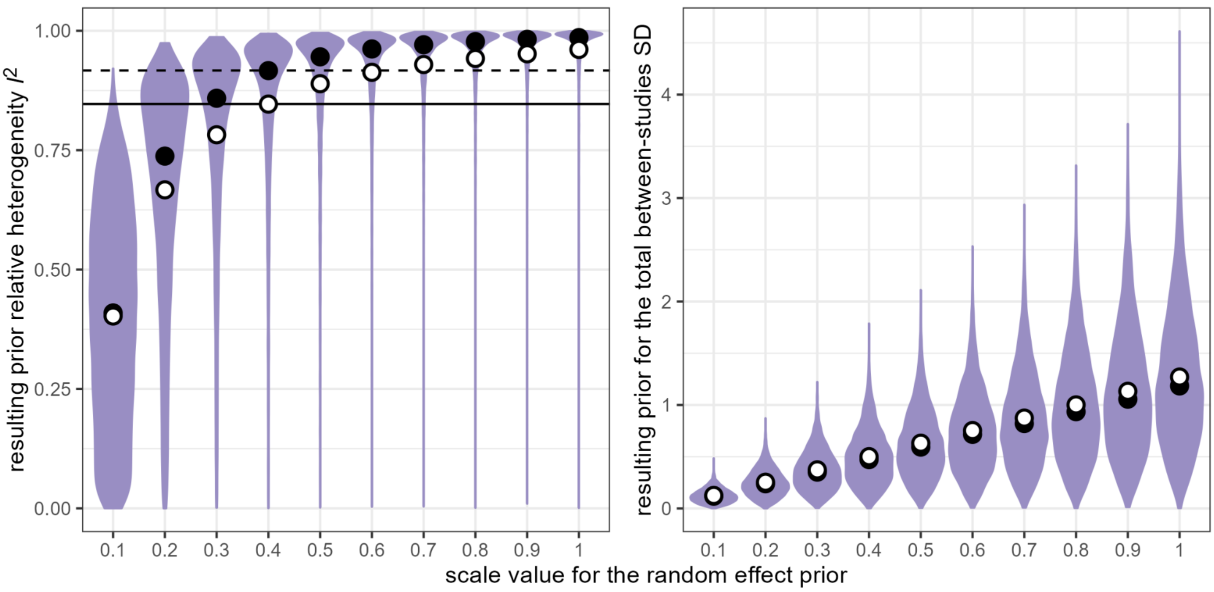


**Figure S4.1.** Left: prior predictive distribution of relative heterogeneity $I^2$ under various scales for the random effect prior. The black dots correspond to the medians, the white dots to the means. The horizontal lines correspond to the median (full line) and mean (dashed line) estimates from Senior et al. (2016) (note the inverse ordering compared to the dots). Right: prior distribution of the total between-studies SD, under various scales for the random effect prior.

#### Supplementary Material 5: Supplementary results

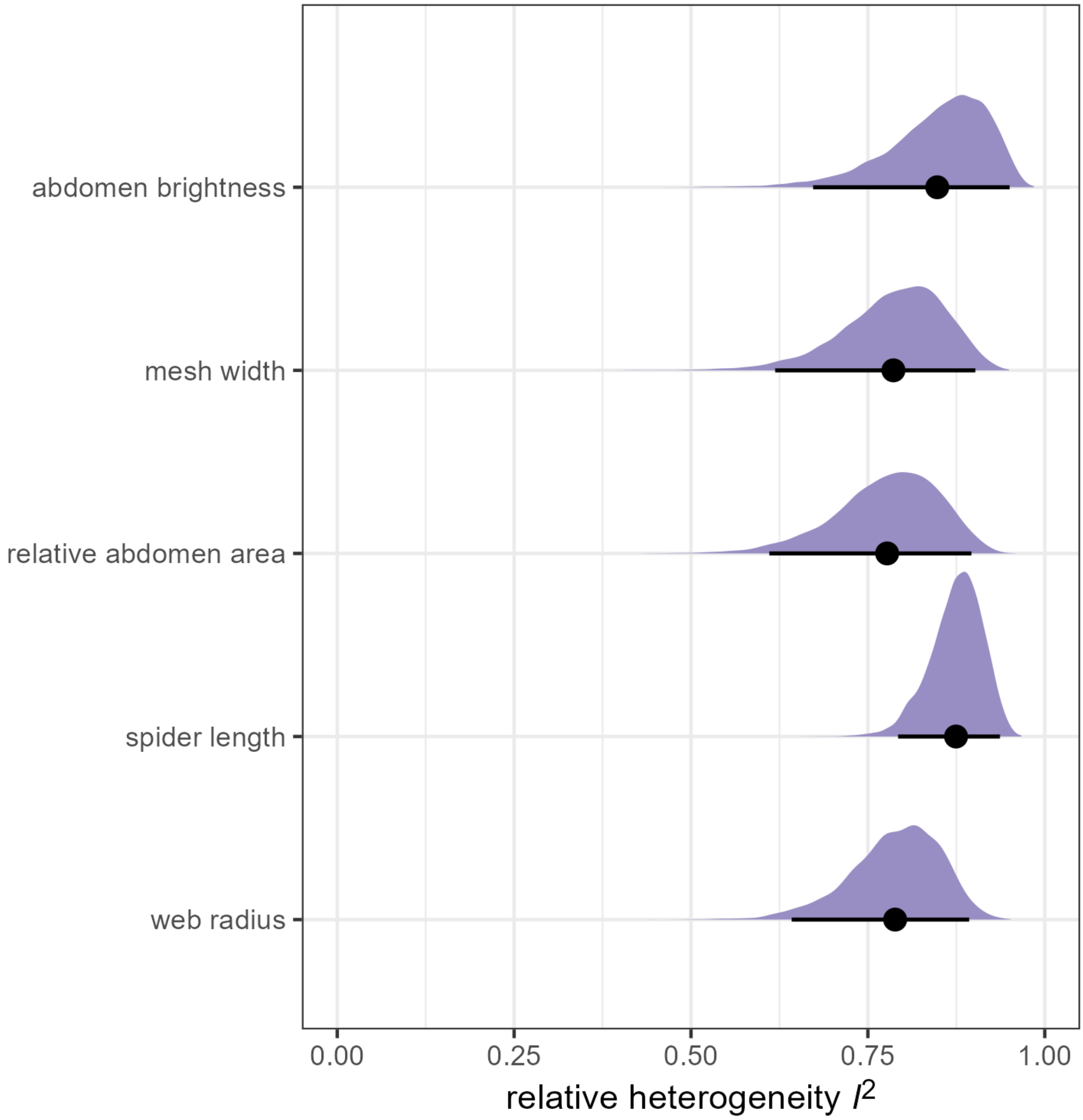


**Figure S5.1.** Posterior distributions of relative heterogeneity estimates $I^2$.

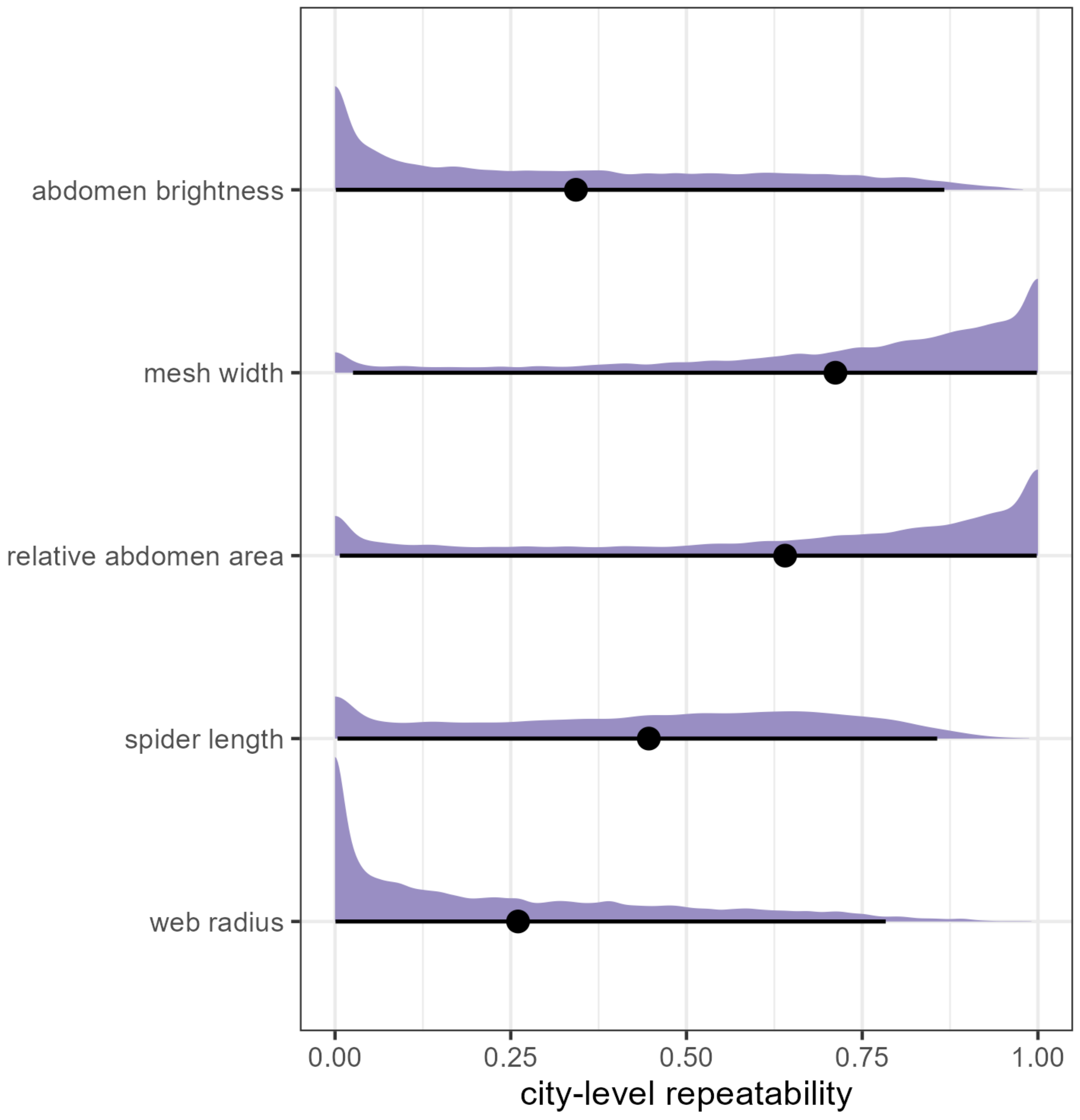


**Figure S5.2.** Posterior distributions of the proportion of absolute heterogeneity $\tau^2 + \sigma^2$ due to the between-cities variance component $\tau^2$

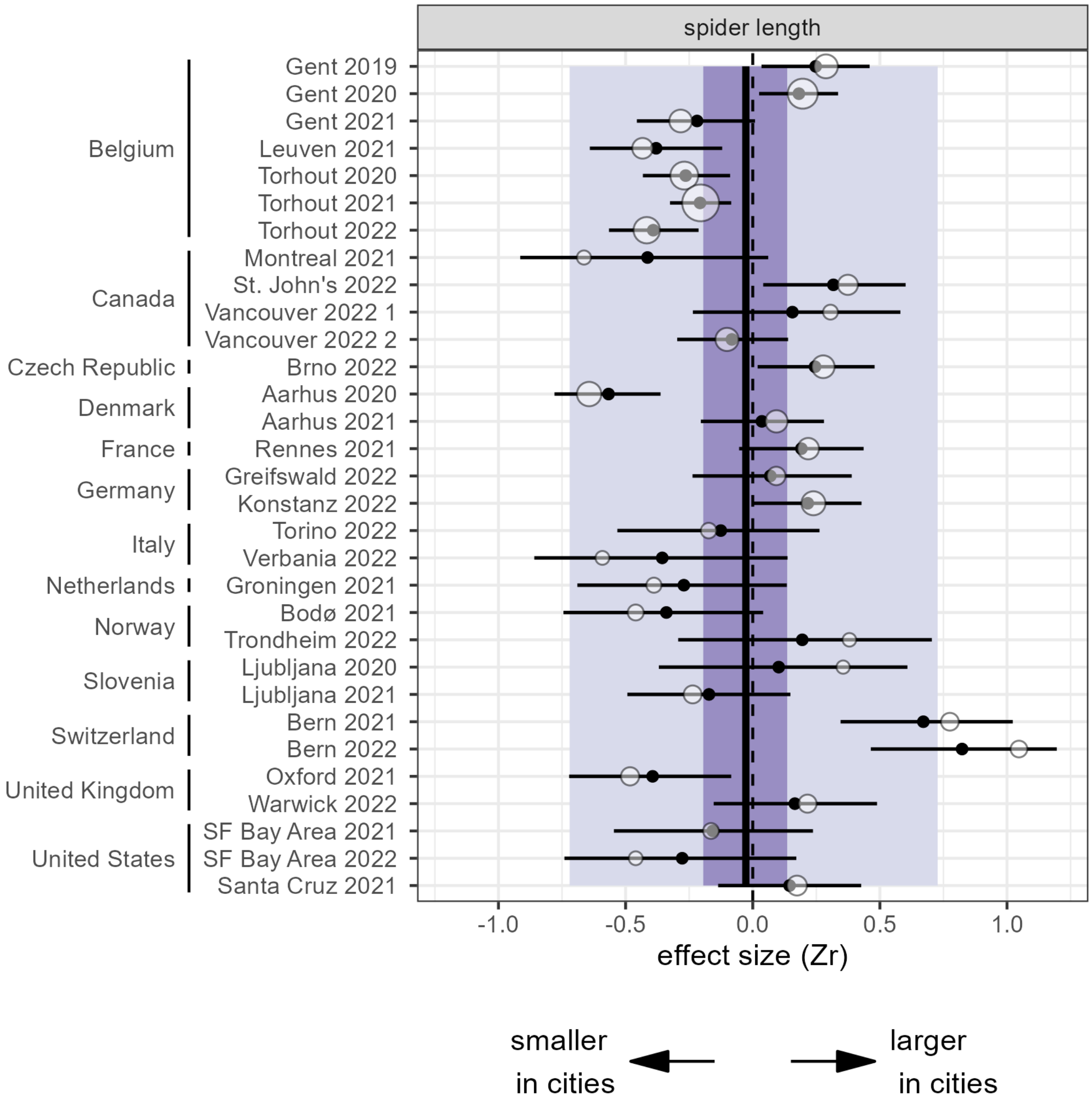


**Figure S5.3.** Observed and predicted effect sizes for spider length, based on the intercept-only model. As in main text **Figure 4**, white bubbles are observed effect sizes with their width inversely proportional to sampling variance, while black dots and segments are predicted effect sizes and their 95% credible intervals from the models. The darker colored bands are the 95% credible intervals for the overall meta-analytic means, while the lighter colored bands are 95% prediction intervals, i.e. indicate the expected distribution of individual effect sizes.

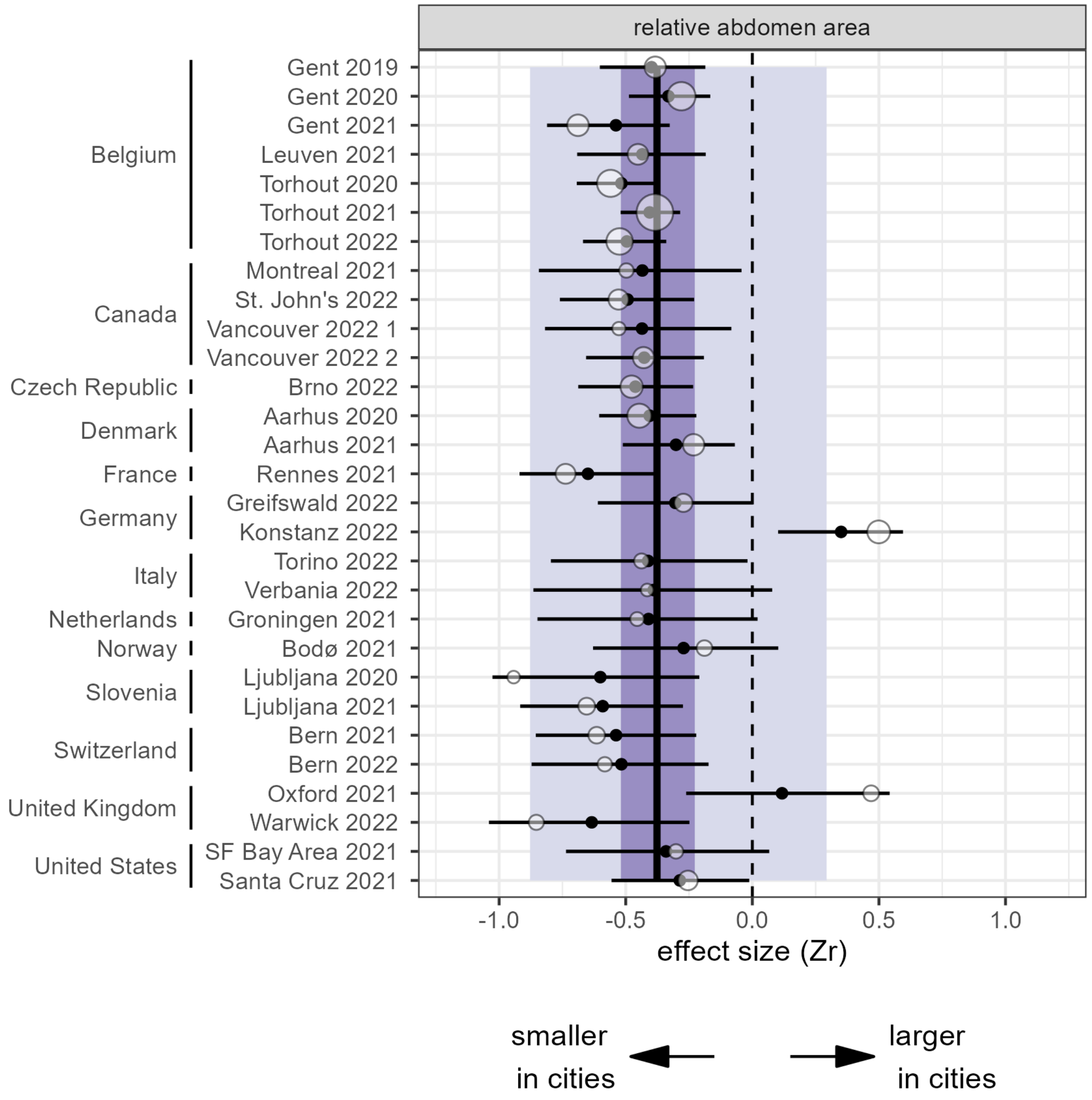


**Figure S5.4.** Observed and predicted effect sizes for relative abdomen area, based on the intercept-only model. Legend is as in **Figure S5.3**.

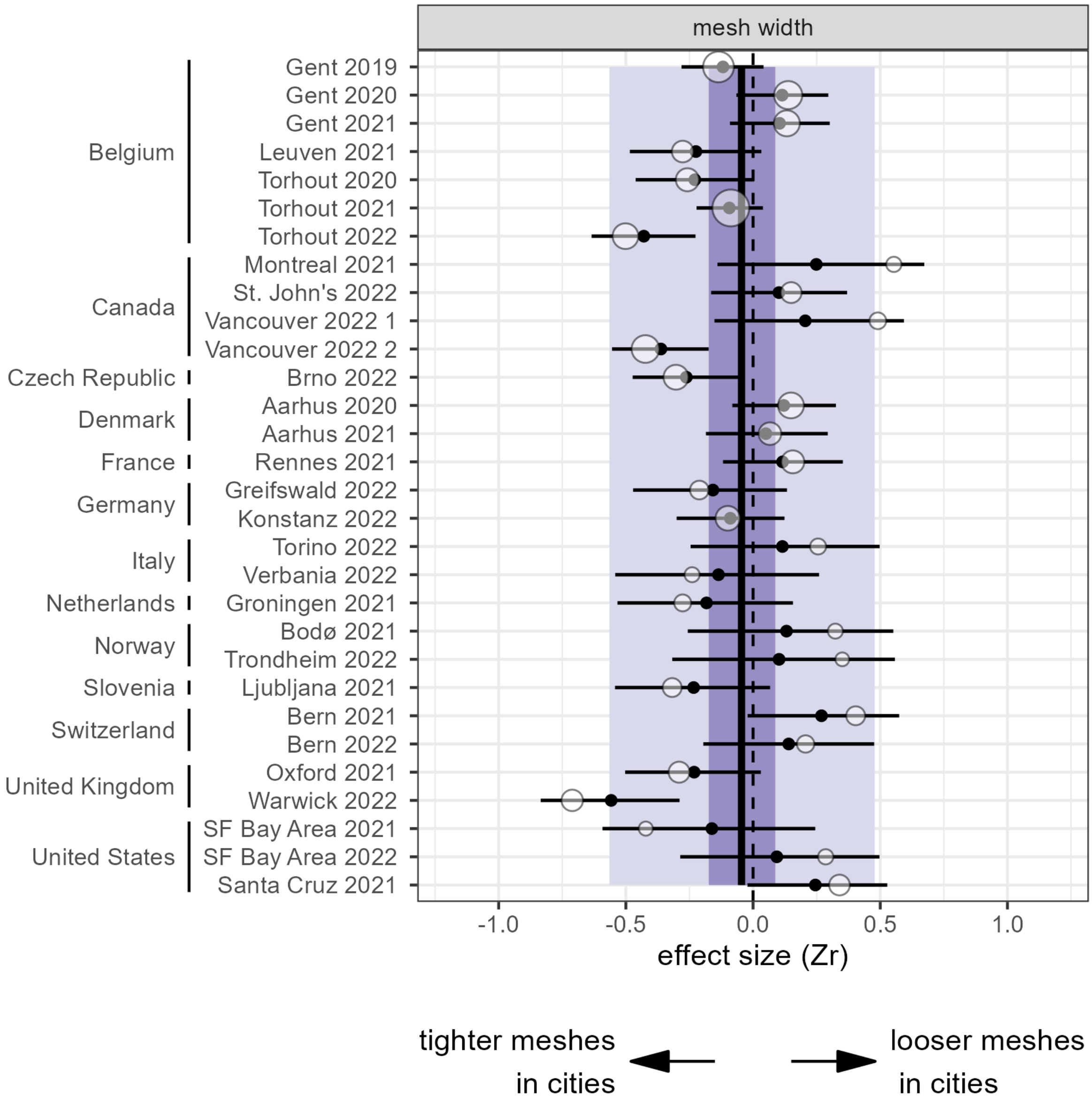


**Figure S5.5.** Observed and predicted effect sizes for web mesh width, based on the intercept-only model. Legend is as in **Figure S5.3**.

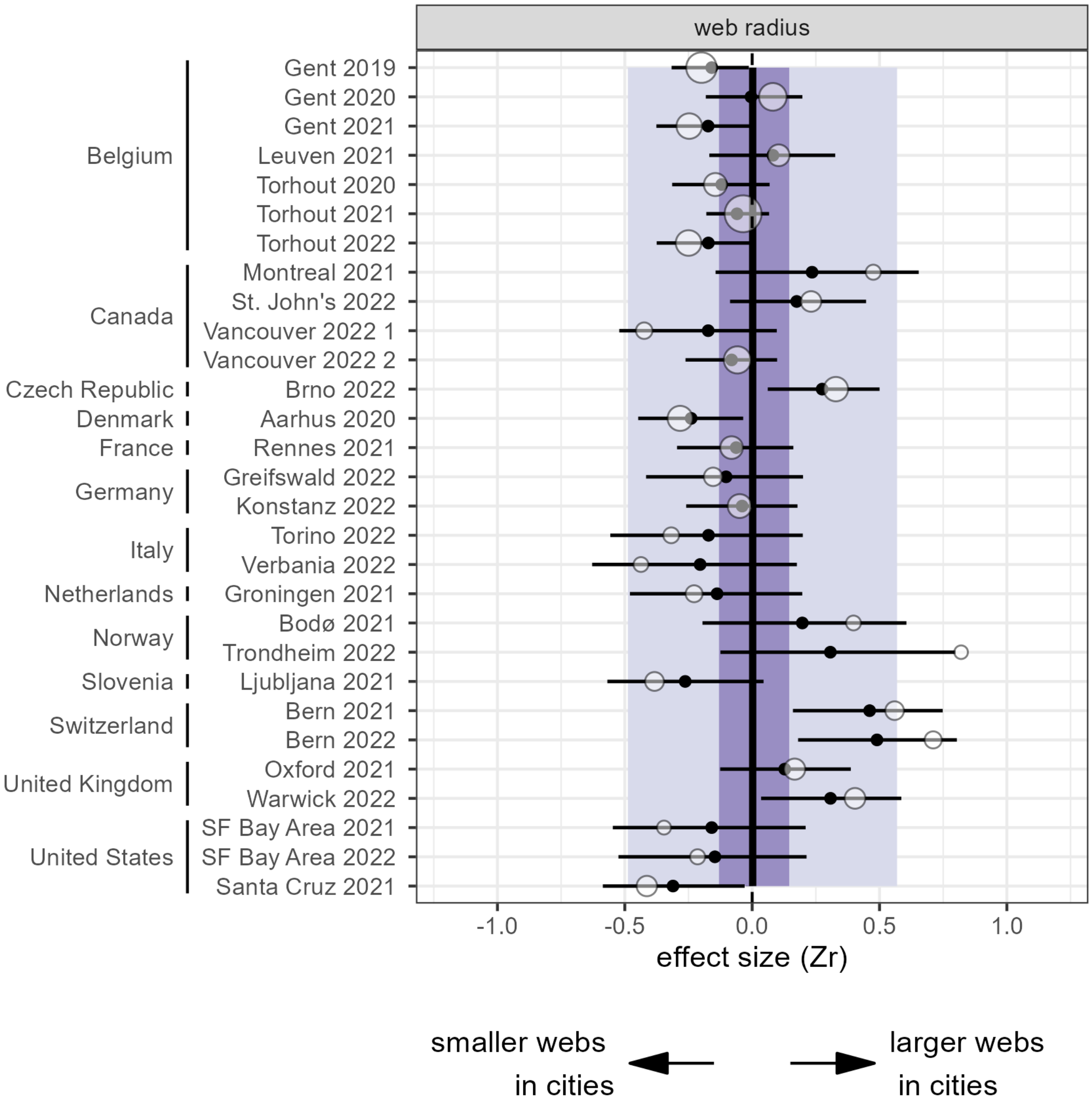


**Figure S5.6.** Observed and predicted effect sizes for web radius, based on the intercept-only model. Legend is as in **Figure S5.3**.

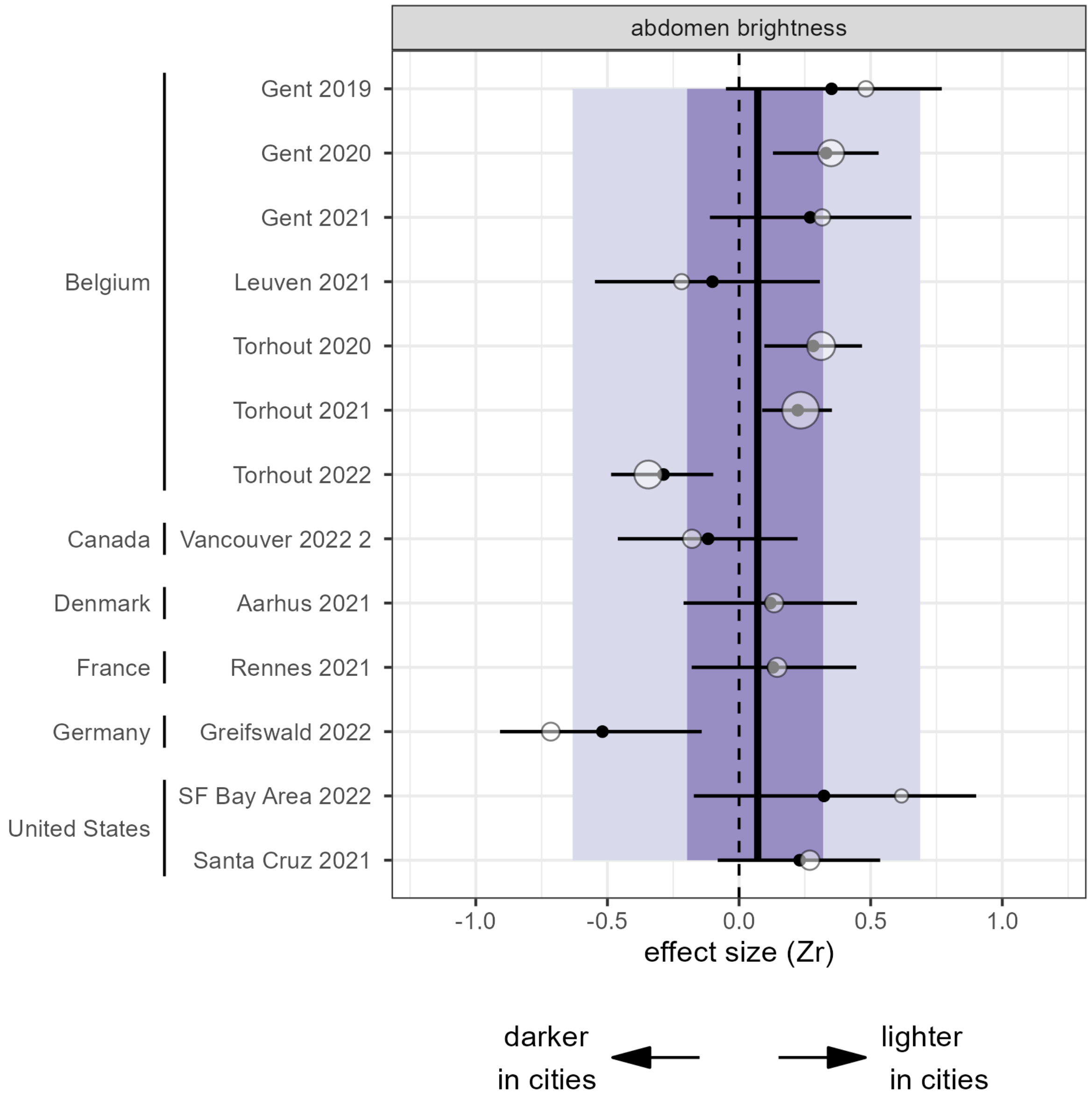


**Figure S5.7.** Observed and predicted effect sizes for abdomen brightness, based on the intercept-only model. Legend is as in **Figure S5.3**.

**Table S5.1.** Summary of moderator coefficients from meta-regressions with all selected moderators included at the same time (compare with main text **Table 2**). No multi-moderator model was fitted for abdomen brightness, given the limited number of effect sizes for that trait (see main text **Methods** and **Results** for details). Moderators, but not effect sizes, were centered and scaled to unit 1 standard deviation before model fitting.

| coefficient | spider length | relative abdomen area | mesh width | web radius |
|---|---|---|---|---|
| Intercept | -0.02 [-0.18, 0.13] | **-0.40 [-0.53, -0.28]** | -0.03 [-0.17, 0.11] | -0.00 [-0.12, 0.12] |
| annual mean temperature | -0.07 [-0.25, 0.11] | 0.06 [-0.08, 0.19] | -0.07 [-0.23, 0.08] | **-0.25 [-0.39, -0.11]** |
| size of urban area | -0.19 [-0.41, 0.04] | 0.06 [-0.13, 0.24] | 0.12 [-0.09, 0.32] | -0.01 [-0.20, 0.18] |
| tree cover | 0.15 [-0.05, 0.34] | 0.02 [-0.14, 0.18] | -0.05 [-0.22, 0.12] | 0.00 [-0.16, 0.15] |
| mean Human Modification index | -0.05 [-0.23, 0.12] | 0.03 [-0.12, 0.16] | -0.07 [-0.23, 0.09] | 0.05 [-0.09, 0.19] |
| mean sampling date | **0.24 [0.06, 0.41]** | **-0.21 [-0.35, -0.05]** | -0.01 [-0.18, 0.15] | 0.07 [-0.09, 0.22] |

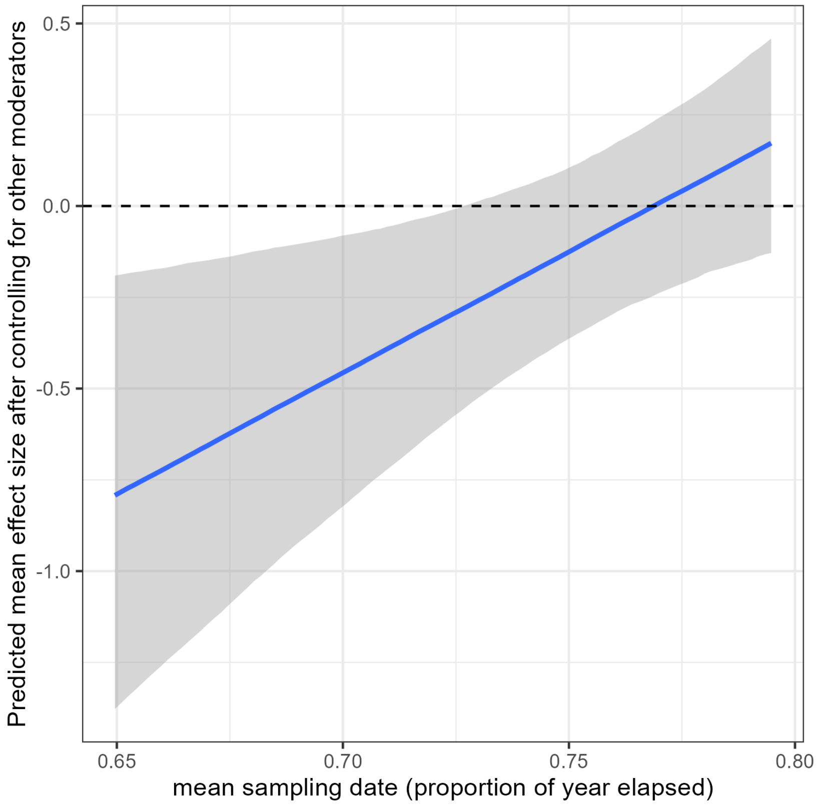


**Figure S5.8.** Mean (and 95% credible interval) conditional effect of sampling date on spider length effect sizes, after accounting for the other moderators (based on the model in **Table S5.1**).
